# Opioid-Induced Inter-regional Dysconnectivity Correlates with Analgesia in Awake Mouse Brains

**DOI:** 10.1101/2024.07.30.604249

**Authors:** Jean-Charles Mariani, Samuel Diebolt, Laurianne Beynac, Renata Santos, Stefan Schulz, Thomas Deffieux, Mickael Tanter, Zsolt Lenkei, Andrea Kliewer

## Abstract

The µ-opioid receptor (MOP) is crucial for both the therapeutic and addictive effects of opioids. Using a multimodal experimental approach, here we combined awake functional ultrasound (fUS) imaging with behavioral and molecular assessments, to examine opioid-induced changes in brain activation and functional connectivity (FC). Morphine, fentanyl, and methadone induce significant dose- and time-dependent reorganization of brain perfusion, oscillations and FC in awake mice. Notably, opioids induce a transient, region-specific hyperperfusion, followed by a consistent MOP-specific dysconnectivity marked by decreased FC of the somatosensory cortex to hippocampal and thalamic regions, alongside increased subcortical and intra-cortical FC. These FC changes temporally correlate with generalized brain MOP activation and analgesia, but not with hypermobility and respiratory depression, suggesting a reorganization of inter-regional FC as a key opioid effect.

## INTRODUCTION

The lack of a precise understanding of drug effects on brain function poses a significant challenge in the development of new drugs that act on the central nervous system. This has hindered the development of improved antipsychotics and highly effective analgesics without detrimental side effects (*1*), particularly in light of the opioid crisis.

Extensive research has shown that cognition and behavior rely on the integration, synchronization, and coordination of large-scale sensory and motor networks. The large-scale organization of neural networks and drug-induced changes in connectivity patterns and activation maps may provide valuable testable hypotheses regarding the neural underpinnings and pharmacological accessibility of brain function and behavior. Brain imaging approaches, particularly functional magnetic resonance imaging (fMRI), can bridge the knowledge gap between cellular and behavioral levels by providing a readout of brain activation and functional connectivity (FC) through the neurovascular coupling. In functional brain imaging, FC is defined by the spatiotemporal correlation of spontaneous fluctuations in perfusion. As such, ‘pharmaco-fMRI’ (phMRI), by measuring drug-induced local changes in the Brain Oxygenation Level Dependent (BOLD) signal, an indirect measure of brain perfusion, has been proposed as an interesting tool for both preclinical and clinical drug development (*2, 3*). However, the preclinical use of phMRI is significantly constrained due to the need for immobilization, often accompanied by some degree of anesthesia, and its relatively low sensitivity, particularly in mice, a prominent preclinical rodent model. Anesthesia use poses specific challenges in analgesia research as it hinders the capture of cortico-thalamic connectivity, a critical component of sensory-motor integration, even with advanced fMRI technology (*4*). Medetomidine, an anesthetic agent commonly used in preclinical fMRI studies, has been observed to interfere with cortico-thalamic connectivity even at low doses, affecting brain regions expressing high levels of beta-2 adrenergic receptors (*5–7*). Notably, a recent pioneering phfMRI investigation of opioid (oxycodone) action on mouse brain FC, also utilizing medetomidine anesthesia (*8*), did not report noteworthy changes in the cortico-thalamic network, which is intriguing for an analgesic agent. Consequently, there is a pressing need for innovative technologies that permit large-scale and highly-resolved network and perfusion imaging in awake mice to fully leverage the potential of brain imaging in neuro-pharmacological research.

Recently, functional ultrasound (fUS) imaging, a novel technique based on ultrafast power Doppler imaging (*9*), has been developed for imaging brain perfusion, activation, and FC in rodents (*10*). Conceptually similar to BOLD MRI, the precise measurement of task-induced or spontaneous (resting-state) changes in cerebral blood volume (CBV) in individual voxels allows to establish brain activation and FC patterns, respectively, through the neurovascular coupling. fUS imaging offers very high sensitivity and spatiotemporal resolution (sub-second time resolution and 100 μm in-plane spatial resolution), with elevated operating simplicity when compared to fMRI methods. By employing fUS imaging, previously unknown facets of brain function can now be explored (*11–13*). We have specifically adapted fUS for imaging task-related responses and FC in awake mice^14^ and our recent proof-of-concept study has demonstrated that fUS imaging can effectively visualize pharmacologically-induced alterations in FC in the awake mouse brain, indicating its promise as an emerging valuable tool for neuropharmacological research (*14*).

In this study, we aimed to investigate whether clinically relevant opioids, acting on the µ-opioid receptor (MOP), elicit consistent and reproducible alterations in regional brain activation and functional connectivity. Specifically, we sought to determine if we could establish a robust, dynamic and pharmacologically relevant fingerprint of opioid action in awake and behaving mice, through the intact skull. To assess the pharmacological relevance of opioid-induced FC alterations, we studied dose- and time-dependence, drug-specific pharmacodynamics, sensitivity to pharmacological and genetic MOP inactivation, and manifestation of habituation/tolerance following chronic treatment. As a pioneering endeavor, our hypothesis centered on the significant changes in FC between neocortical, hippocampal, and thalamic regions, achieved through recording on a single, easily localized coronal brain slice encompassing these specific regions. We enhanced the sensitivity of our functional fingerprinting by employing a data-driven approach to define individual functional regions-of-interest (ROIs) within these brain areas. Finally, after establishing the pharmacological relevance of the observed FC alterations, we have correlated changes in FC with specific CBV time series, behavioral (locomotion, analgesia, respiration) and molecular (MOP phosphorylation as a proxy measure of MOP activation) readouts. Together, our results validate a specific fingerprint for a drug family with high medical and societal impact. These results also lay the foundation for a new approach in the development of highly-effective pain relievers and other neuropsychiatric drugs. This new method has the potential to identify compounds with improved pharmacological profiles, through systematic and quantitative screening of drugs effect on the FC network.

## RESULTS

### Pharmacological fUS fingerprint of morphine in awake mice

To investigate pharmacologically-induced changes in brain activation and FC in awake and behaving mice, we adapted the imaging protocol from our previous work (*15*). Following implantation of a small metal headplate on the mouse skull and several days of habituation, the mouse is head-fixed within the Mobile HomeCage®, an air-lifted lightweight carbon cage facilitating continuous tracking of the animal behavior (*16*), during imaging experiments (Fig. 1A-C). Our standard coronal imaging plane was selected at Bregma -1.8 mm, encompassing key regions of interest, such as the somatosensory cortex, the retrosplenial area, the dorsal hippocampus and the thalamus (Fig. 1D-H and Fig. S1). The choice of this specific coronal slice was based on its inclusion of major regions of interest and its distinct display of a readily identifiable branching configuration between the anterior choroidal and the internal carotid arteries (Fig. 1E), facilitating reproducible localization within and between mouse cohorts (Fig. S1A and B). The imaging plane (Fig. 1D) was perpendicular to the metal headplate, exhibiting an 8° angle in the coronal plane with respect to the Paxinos mouse brain atlas (*17*) and an 18° angle with respect to the Allen mouse brain atlas (*18*). All images were registered to a custom vascular template of the coronal slice of interest. Our treatment protocol (Fig. 1G) involved a 20 min baseline period divided into two 10-minute sub-phases (BP 1 – BP 2), followed by a 60 min post-injection "drug phase" period, further divided into 10 min sub-phases (SP 1 – SP 6). To enhance the sensitivity of our functional fingerprinting, we defined 18 unilateral and symmetrical functional regions-of-interests (ROIs) by clustering a voxel-wise correlation matrix (see Methods) (Fig. 1H and Fig. S2).

**Fig. 1.**
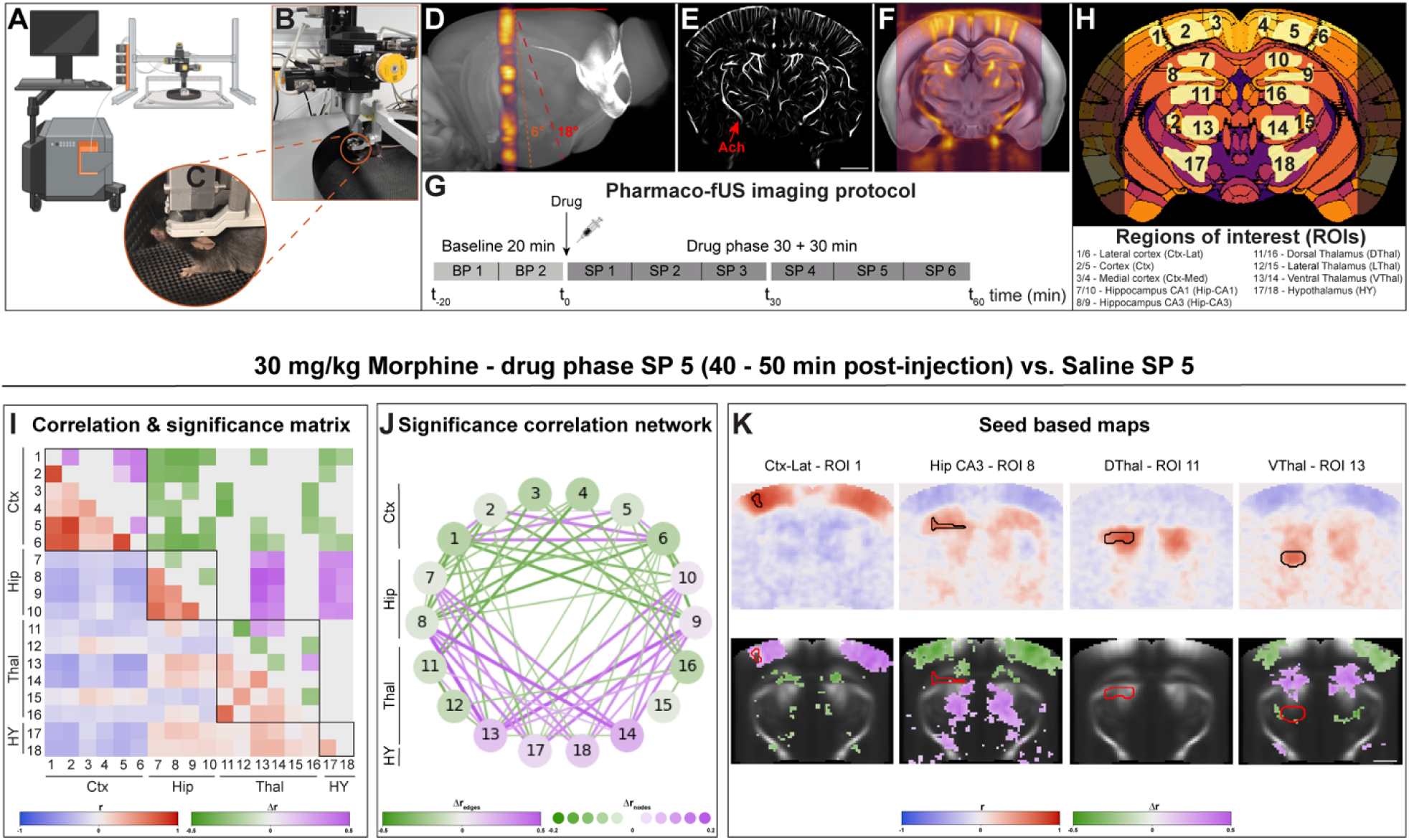
Pharmaco-fUS imaging in awake mice. (**A**) Schematic representation of the fUS imaging setup with the Mobile HomeCage® setup. (**B** and **C**) Transcranial acquisition of a head-fixed and freely moving mouse through a chronically implanted metal frame with an ultrasonic probe. (**D**) Localization of the imaged plane. The solid red line represents the plane of the metal plate, while dashed orange and red lines represent the coronal plane in the Paxinos Mouse Brain Atlas (6° difference) and Allen Mouse Brain Common Coordinate Framework (CCFv3) reference space (18° difference), respectively. (**E**) Super resolution image of the imaged plane with 25 μm resolution of the vascular tree. The anterior choroidal artery (Ach, red arrow) was used for reproducible localization of our reference imaging plane. (**F**) Superimposition of the imaging plane from a single animal with the Allen brain template. (**G**) Pharmaco-fUS imaging during the acute treatment protocol. Three recording periods (one baseline, and two drug phases) are split in 10 min windows (baseline-phase: BP 1-2 and sup-phase: SP 1-6, respectively). (**H**) Selected regions of interests (ROIs) in the coronal plane. Data-driven parcellation in the imaged plane superimposed with the Allen brain parcellation. 18 ROIs identified by the algorithm and having a good match with major anatomical regions were used. (**I** and **K**) Averaged analysis of the FC pattern after injection of 30 mg/kg morphine in wildtype (C57BL/6J) mice (n = 8 – 10) compared to control (saline injection), at 40 – 50 min post-injection (SP 5). (**I**) ROIs based functional connectivity correlation/significance matrix. Numbered rows and columns represent ROIs from Fig. 1H. Squares delineate four meta-regions (cortex, hippocampus, thalamus and hypothalamus). Bottom left triangle shows the average post-injection correlation matrix in blue-red ([−1, 1]), top right triangle displays the effect size (difference of correlation) in green-violet ([−0.5, 0.5]) of ROI pairs where difference with the saline state is significant. (**J**) Significance correlation graphs representing the significance part of the correlation matrix where the nodes encode the global connectivity of each ROI in the network, and edges the significant changes in their correlation in green-violet (edges [−0.5, 0.5]; nodes [−0.2, 0.2]). (**K**) Seed-based maps of the ROIs 1, 8, 11 and 13. Top row represent the difference average correlation map in blue-red ([−1, 1]) and bottom row the voxels, whose difference with saline injected animals is significant, are overlaid on the PD template, the size effect is displayed in green-violet ([−0.5, 0.5]). Scale bar = 1 mm.

Following the s.c. administration of 30 mg/kg morphine, a strong and statistically significant alteration in averaged brain FC patterns (n = 8 – 10) was observed when compared with the saline-treated control cohort (n = 8 – 10). At 40 min post-injection, a reduction in cortical-subcortical FC coincided with increased FC within cortical regions and increased hippocampo-thalamic FC, as depicted in the connectivity/significance matrix (Fig. 1I), the significance correlation network (Fig. 1J), the seed-based maps (Fig. 1K), and quantified in key ROI pairs (Fig. S3). This result indicates that morphine induces a significant redistribution of both intra- and inter-regional FC patterns in the awake mouse brain, with a prominent effect characterized by the emergence of an anti-correlation between cortical and subcortical regions.

Next, we tested the pharmacological relevance of our findings. As expected, injection of saline did not yield any noticeable changes in FC during the entire acquisition period (Fig. 2A and B). In contrast, increasing doses of morphine resulted in a progressively earlier onset of the morphine-induced reorganization of the functional connectome, likely indicating an earlier attainment of the critical level of MOP receptor occupation, with higher morphine doses leading to stronger FC changes (Fig. 2A and B, Fig. S3). Strikingly, even the highest administered dose of morphine (70 mg/kg) failed to induce similar connectivity changes in MOP^-/-^ knockout mice (Fig. 2C-D). Similarly, pre-treatment with naloxone, a fast-acting MOP-specific antagonist, significantly attenuated the effects of 30 mg/kg morphine at 20-30 minutes (SP 3) after injection (Fig. 2A and B).

**Fig. 2.**
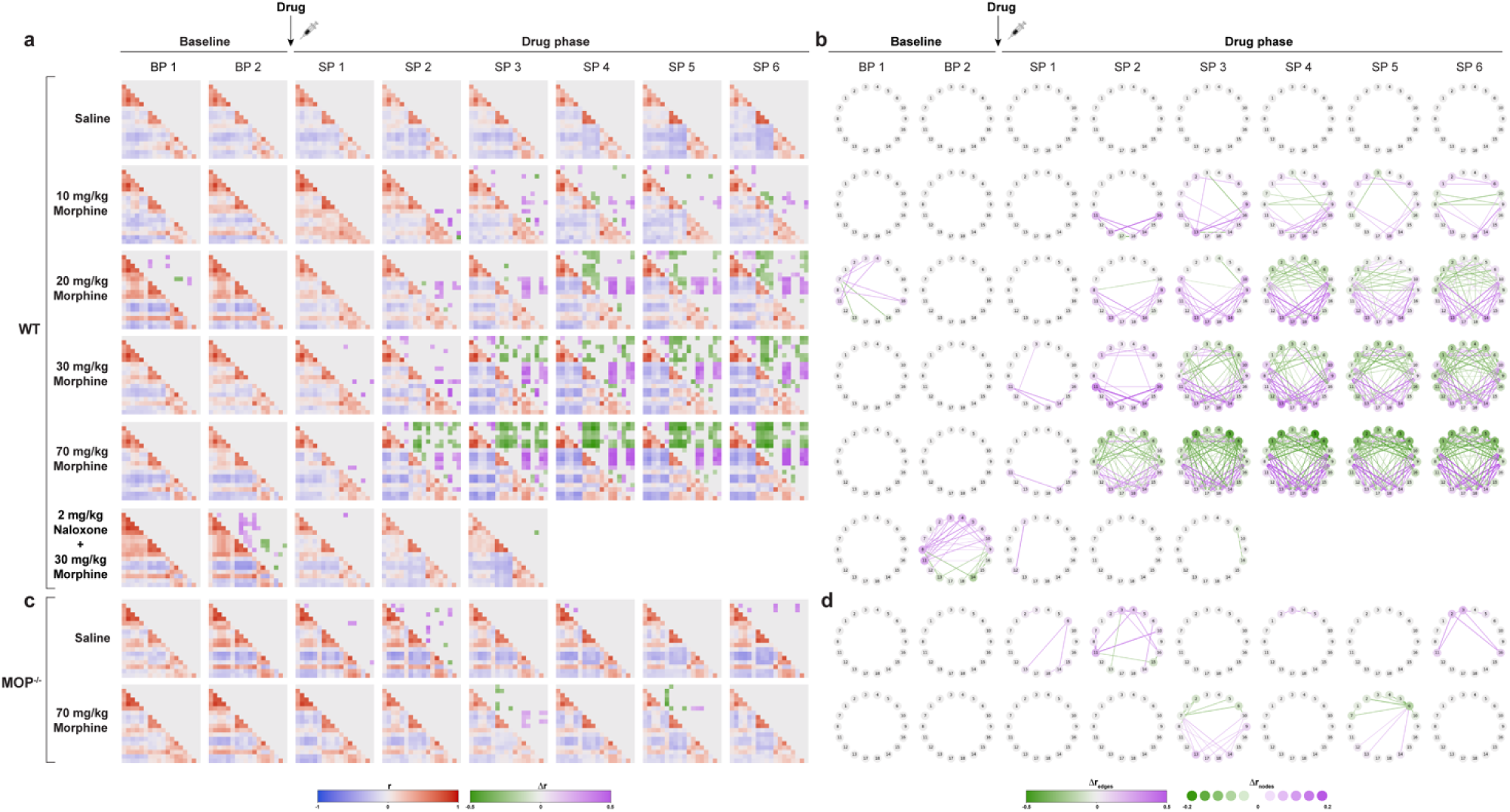
Dose- and time-dependence of FC signatures of morphine. (**A – C**) Pharmacodynamics and dose dependence of the morphine FC pattern in (**A** and **B**) wildtype (C57BL/6J) or (**C** and **D**) MOP knockout mice (MOP^-/-^) (n = 8 – 10). Naloxone pre-treatment was at t_-5_ minutes, followed by morphine at t_0_. FC correlation/significance matrices and significance correlation graphs are shown as on Fig. 1I and J. Statistical significance of the morphine FC pattern by comparing drug phase to saline injection. Correlation matrix in blue-red ([−1, 1]); significance matrix in green-violet ([−0.5, 0.5]); significance correlation graph in green-violet (edges [−0.5, 0.5]; nodes [−0.2, 0.2]).

### Temporal dynamics of the morphine fingerprint

In order to gain mechanistic insight into morphine-induced changes in FC, we analyzed the spatio-temporal reorganization of the power Doppler signal patterns. In control animals, the power density of the power Doppler signal in the frequency domain displayed an inverse correlation with frequency in distinct ROIs, consistent with a power law distribution (Fig. 3A). Similar 1/f statistical patterns are expected to arise from the integration of a large family of neuronal oscillations in the spatial and temporal domains (*19*). According to the prevailing consensus, larger neuronal assemblies are associated with slower oscillations characterized by higher amplitudes, while smaller, localized clusters of neurons contribute to higher frequency signals characterized by lower amplitudes (*19*). Intriguingly, a disruption in this power-law relationship in the power Doppler spectrum can be observed starting from 20 minutes after morphine administration: the power of low frequency components between 0.01 and 0.1 Hz is notably redistributed (Fig. 3A). This shift suggests that morphine treatment induces alterations in large-scale FC patterns through perturbation of specific oscillatory components from large neuronal assemblies that contribute to the critical state at rest (*20*). This perturbation of hemodynamic spectral content can be further investigated by seed-based cross-correlation analysis (Fig. 3B and Fig. S4 and Video S1) which confirmed notable changes in large-scale oscillation patterns, as compared to the saline control (Fig 3C-D). Interestingly, contralateral (i. e. intra-regional) ROIs displayed mostly temporally symmetrical cross-correlations in the cortex, hippocampus and thalamus, which were mostly preserved despite of increased oscillation frequencies. In contrast, several ROI pairs spanning different regions showed asymmetrical lagged correlation structures in controls, which were significantly perturbed after morphine treatment, putatively related to altered signal propagation characteristics. Notably, several such inter-regional ROI pairs that did not show significant FC changes at the classical zero lag correlation, such as the lateral cortex-dorsal thalamus connection (ROI 1 to ROI 11), displayed significant FC changes at a lags of 8 – 10 seconds, suggesting that morphine perturbs signal propagation characteristics also in these inter-regional connections (Fig. S4).

**Figure 3.**
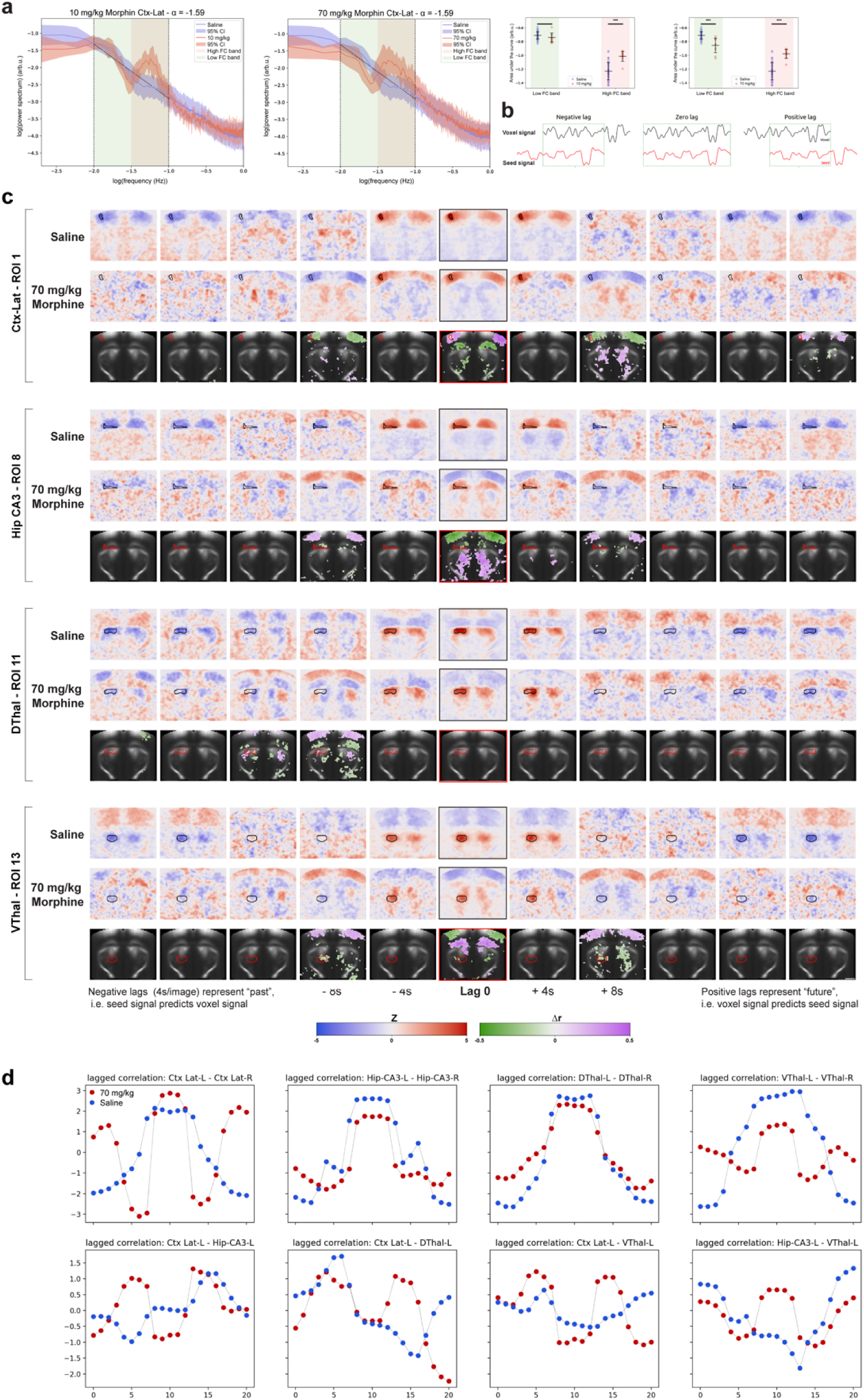
Dynamic effects of morphine on FC. (**A**) Power spectrum analysis (Welch method, normalized integral per animal, 95% CI) in the lateral cortex (ROI 1) following injection of saline (n = 47) or morphine (10 and 70 mg/kg, n = 8 – 10), presented in a log-log diagram. For saline, a characteristic 1/f (i. e. power law) relationship is observed in the FC band (blue line, linear fit dotted black, slope = −1.59). Injections of 10 or 70 mg/kg morphine disrupt the power law relationship and a peak emerges around 0.05Hz (red line), revealing a significant reorganization between low and high FC bands of cortical activity associated with an oscillatory pattern. (**B** and **C**) Spatial analysis of FC dynamics. (**B**) The seed signal is represented in red, while signals from all other voxels are shown in black. Left: Negative lag represents a time shift wherein positive correlation indicates that the seed signal correlates with the past history of the voxel. Middle: Zero lag, which corresponds to classical static FC as shown systemically in our study. Positive correlation indicates that both signals are in phase, Right: Positive lag corresponds to a time shift where positive correlation indicates that future history of the signal correlates with the seed. (**C**) The effects of 70 mg/kg morphine (n = 10) in SP 5 is examined over time, with each window separated by 4 seconds and with seeds in the lateral cortex (ROI 1), hippocampal CA3 (ROI 8), dorsal thalamus (ROI 11) and ventral thalamus (ROI 13). Upper rows: Lag analysis after saline injection, connectivity in each ROI is predominantly bilateral and modulated by a slow oscillation at 30 – 40s periods. Middle rows: After morphine injection the cortical oscillation frequency increases (∼20s period) and the FC pattern becomes more complex with the emergence of two anticorrelated networks. Lower rows: Seed based maps of the significant difference between both conditions. Notably, most ROIs show significant effect at all phases of the fast oscillation induced by morphine, but the dorsal thalamus shows no significant difference at zero lag but a spatial reorganization at the negative lag of -8 – -10 seconds (see also Fig. 1D). See Supplementary Videos for dynamic illustration of these data. Seed based maps in blue-red ([−5, 5]); seed based maps significance in green-violet ([−0.5, 0.5]). Scale bar = 1 mm. (**D**) Inter-regional correlation across lags, each figure represents a pair of ROIs (seed of the SBM and where the correlation is measured: average of the lagged map in the ROI as z-score). The cortical oscillation at a fast frequency is the main driver of temporal reorganization. If some ROI pares show temporal symmetry (top line), hippocampal and thalamic asymmetry, suggesting propagatory phenomena are perturbed by the fast oscillation.

These findings collectively indicate that morphine elicits a MOP-specific, dose- and time-dependent impact on region-wide FC patterns, through the modification of intra- and inter-regional oscillatory behavior, generating altered standing and propagating FC waves. Our results also show that while comprehensive pharmacological FC fingerprinting should extend to the temporal dynamics domain, the classical zero-lag Pearson’s correlation measurements sample already well large-scale changes in FC patterns, allowing us to focus on this approach in the following analyzes.

### Acute and chronic FC fingerprints of major opioid ligands

To further explore whether the MOP-specific effects observed with morphine extend to other MOP-acting ligands, we investigated the brain effects of injecting fentanyl, methadone or buprenorphine, three therapeutically important drugs (Fig. 4). The dose-dependent effects of fentanyl, a rapidly-acting MOP agonist, were measured, along with single doses of 10 mg/kg for methadone and 3 mg/kg for buprenorphine. Similar to morphine, increasing doses of fentanyl elicited increasingly earlier and more pronounced changes in FC, displaying a comparable interregional dysconnectivity pattern (Fig. 4 and Fig. S5 and S6). Similar pattern emerged after administration of methadone, a full MOP agonist and, with a decreased intensity, after administration of buprenorphine, a partial MOP agonist, collectively showing that intensity differences in the change of FC patterns are correlated with differences in MOP agonist efficacy (Fig. S6).

**Fig. 4.**
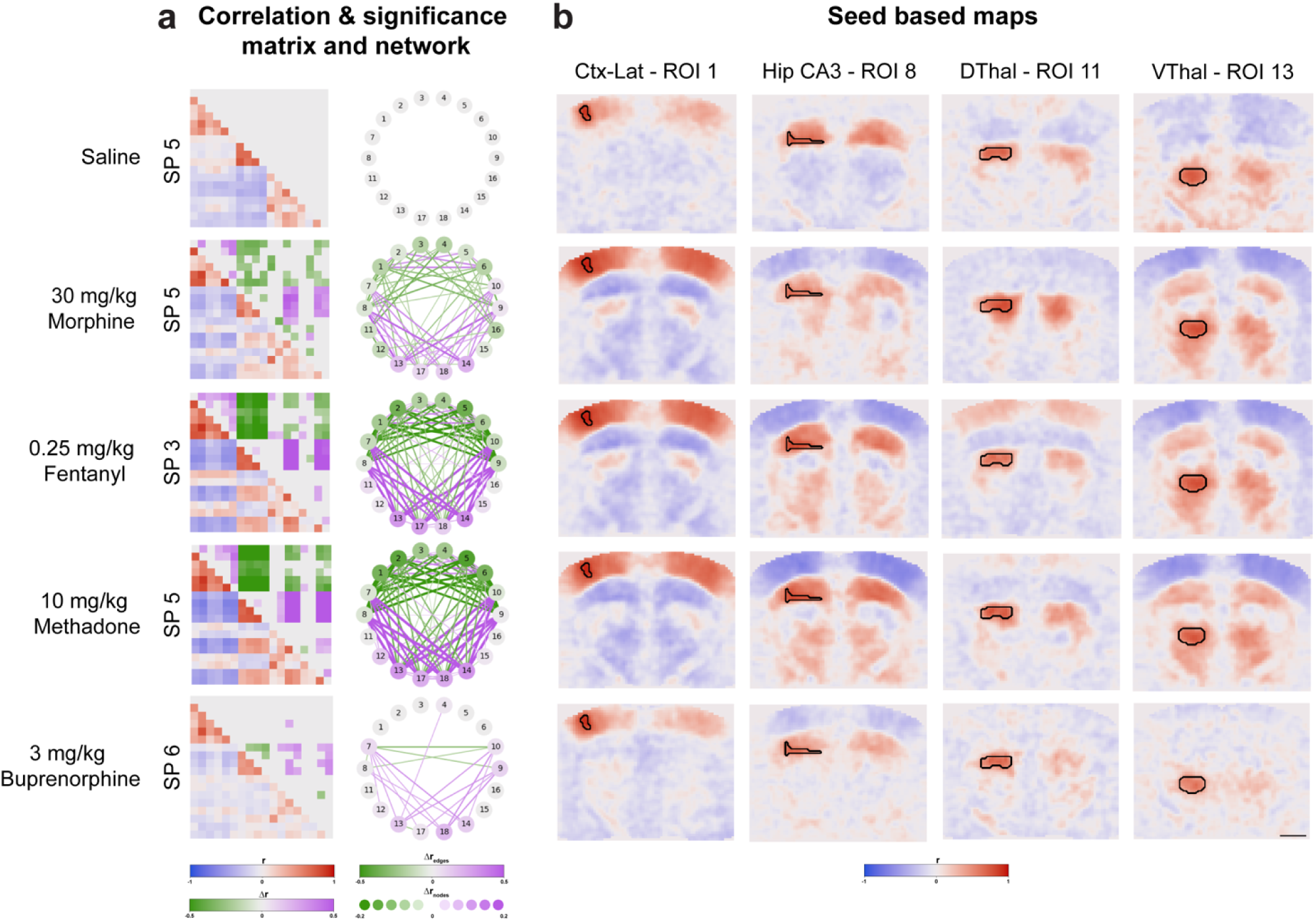
FC signatures of different opioids. (**A**) Functional connectivity correlation/significance matrices and significance correlation graphs at their maximal post-injection effect time points for saline (SP 5), morphine (SP 5), fentanyl (SP 3), methadone (SP 5) or buprenorphine (SP 6) in wildtype (C57BL/6J) mice (n = 8 − 10). (**B**) Seed based maps of ROI 1, 8, 11 and 13. Statistical significance of the FC pattern by comparing drug phase to saline injection. Correlation matrix in blue-red ([−1, 1]); significance matrix in green-violet ([−0.5, 0.5]); significance correlation graph in green-violet (edges [−0.5, 0.5]; nodes [−0.2, 0.2]); seed based maps in blue-red ([−1, 1]). Scale bar = 1 mm.

The behavioral effects of opioids, including analgesia, euphoria, and sedation show tolerance with repeated administration. As a result, patients or drug consumers require higher doses over time to achieve the same level of response, leading to increased risk of side-effects like respiratory depression, constipation, addiction and overdose. We aimed to investigate whether morphine and fentanyl induced changes in FC patterns also display a tolerance effect. We chronically treated mice using an implanted osmotic pump with morphine and fentanyl for one week, which results in significant habituation for analgesia, as assessed by the hot plate test (Fig. S7B) (*21*). Notably, this protocol led to a reduction in significant changes in FC patterns following a single injection, compared to the injection of the same dose in non-chronically treated animals (Fig. 5). These results provide compelling evidence for the functional relevance of the opioid-induced and MOP-specific FC fingerprint, further highlighting its significance in response to different MOP-acting ligands.

**Fig. 5.**
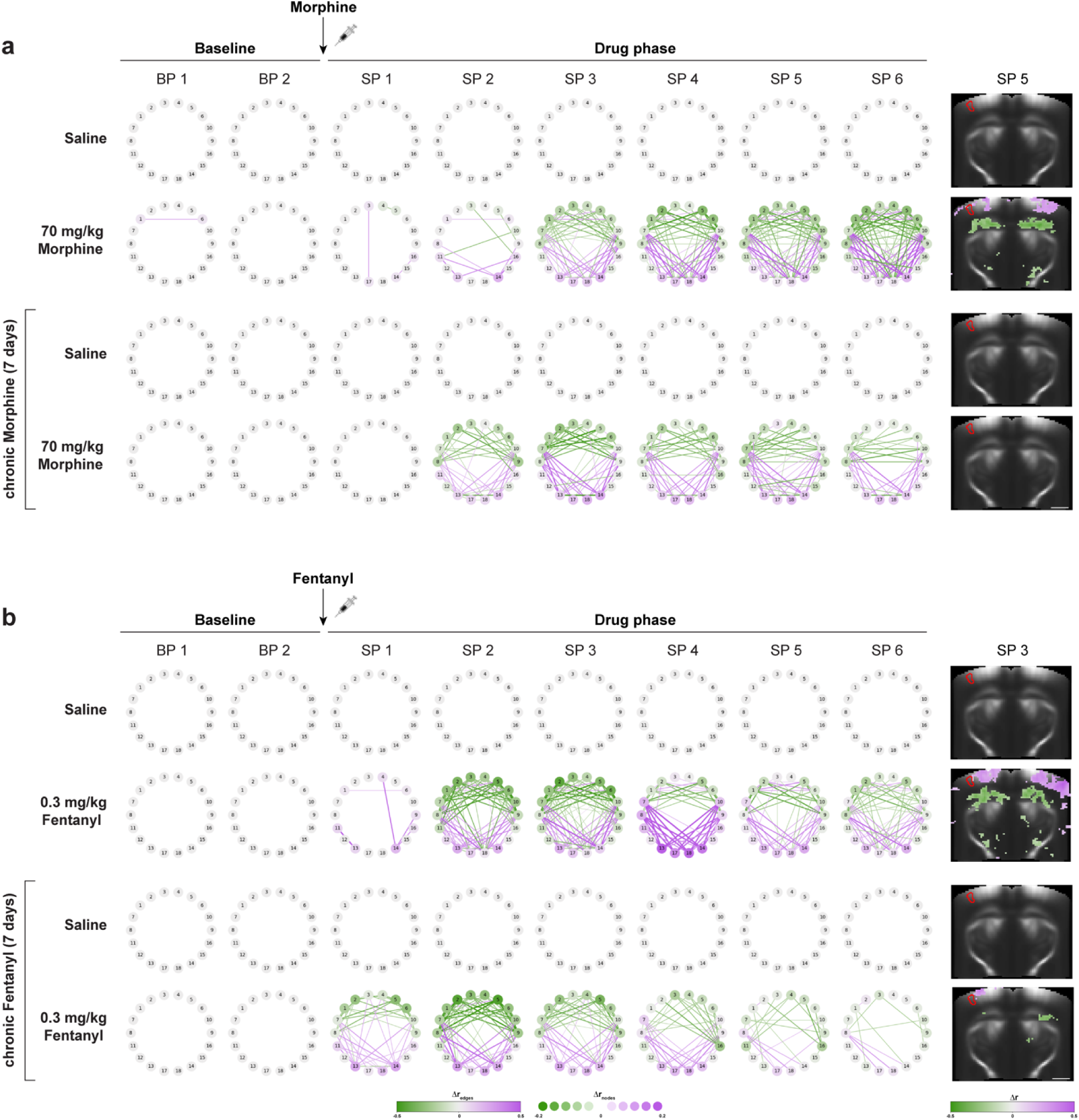
FC patterns of opioid tolerance. (**A – D**) Significance correlation graphs and seed based maps of the lateral cortex (ROI 1 with SP 5 for morphine and SP 3 for fentanyl) after acute saline, morphine or fentanyl injection in wildtype (C57BL/6J) mice (n = 8 − 10). Chronic treatment was carried out with osmotic pumps delivering (**A**) morphine (17 mg/kg/day) or (**B**) fentanyl (2 mg/kg/day) for one week. Statistical significance of the morphine FC pattern by comparing drug phase to saline injection. Significance correlation graph in green-violet (edges [−0.5, 0.5]; nodes [−0.2, 0.2]); seed based maps significance in green-violet ([−0.5, 0.5]). Scale bar = 1 mm.

### Correlation of the FC fingerprint with CBV, behavior and MOP activation

Next, we aimed to understand the relation of the above reported MOP-specific changes in FC to local changes in brain perfusion, the parameter typically measured in ph-fMRI studies, and to opioid-induced behavior. We investigated the correlation of opioid-induced FC changes with alterations in brain perfusion and behavioral read-outs, such as mobility, analgesia and respiratory depression, the latter two measured in parallel awake mouse cohorts (Fig. 6 and Fig. S8). For this purpose, by using principal component analysis (PCA), we derived a FC index to represent with a single parameter the opioid-induced connectome perturbation, i.e. the opioid fingerprint (Fig. S9 and S10). Analysis of CBV changes following injection of morphine or fentanyl, but not of saline, indicated a characteristic change in perfusion (Fig. 6 and Fig. S8). This consisted in early and regionally heterogeneous hyperperfusion followed by prolonged hypoperfusion, with amplitude and dynamics well correlated with the dose and the MOP agonist efficacy of these two ligands. Opioid injection induces a stereotyped locomotion behavior in C57BL/6J but not in BALB/cJ mice (Fig. S11) (*22*). Morphine injection in our C57BL/6J cohorts led to the expected dose-dependent increased mobility (Fig. 6A), which was temporally correlated with changes in brain perfusion. Notably, the morphine-induced patterns of hyperperfusion, albeit with lower intensity, were also present in BALB/cJ mice in the absence of movement, suggesting that regional CBV changes are not a result of increased locomotion (Fig. S11), although they might be amplified by elevated motion (*23*). Finally, morphine induced respiratory depression, a clinically important side-effect of opioids, was also temporally correlated with changes in CBV and locomotion (Fig. 6B).

**Fig. 6.**
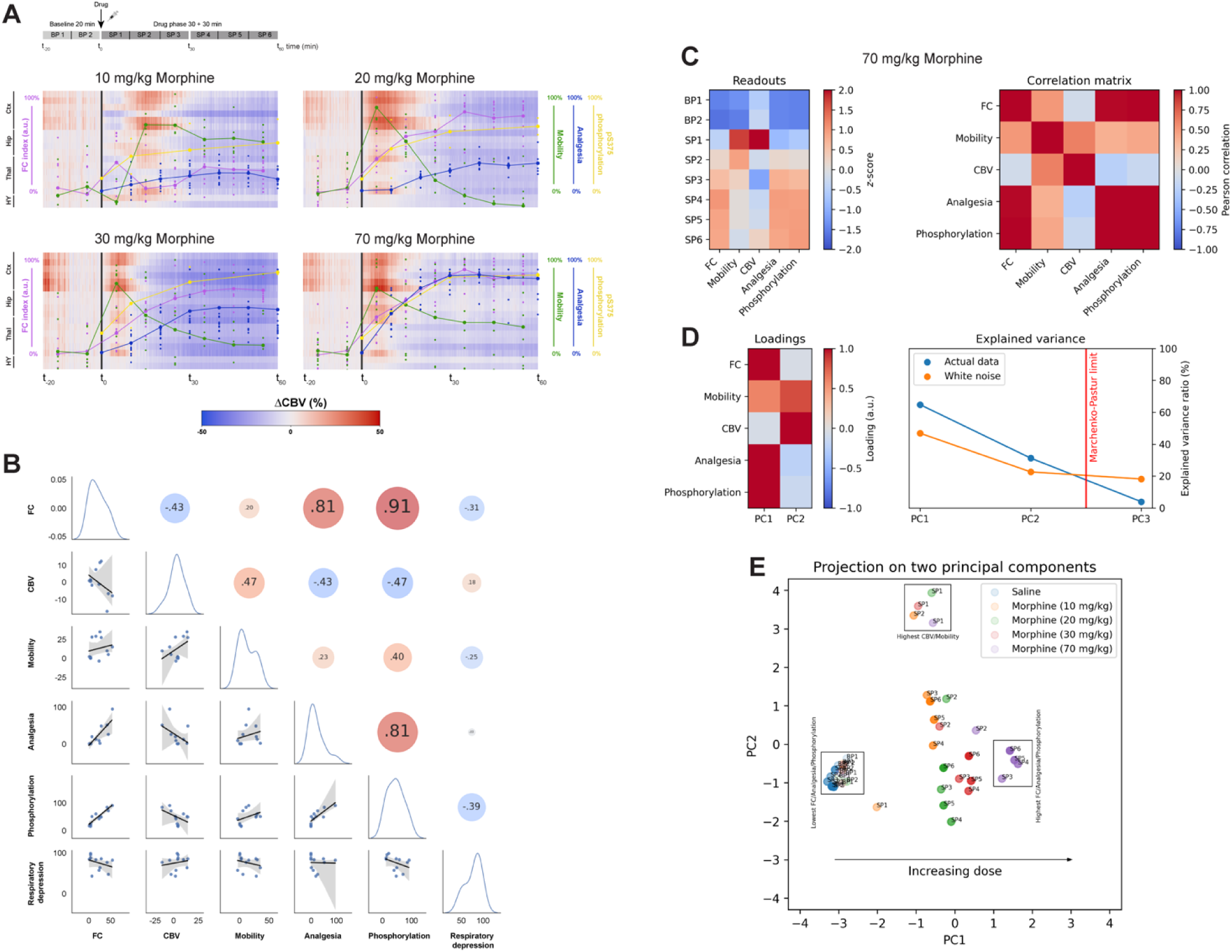
Correlation of morphine-induced FC changes with perfusion, MOP activation and behavior. (**A**) Superimposition of functional connectivity changes (FC index in violet), averaged mobility (green), analgesic response measured in the hot plate test (blue), pS375-MOP phosphorylation pattern detected with Western Blot (yellow) on top of regional CBV change in blue-red ([−50, 50]) after injection of saline and different doses of morphine in wildtype (C57BL/6J) mice (n = 8 − 10). (**B**) Pair plots of correlations between FC, rCBV, mobility, analgesia, MOP phosphorylation and respiratory depression. Graphs in the lower left triangle show correlation between values, each blue dot is the cohort average value for one dose over a 10 min window; black bar is the linear fit and gray area represents the 95% confidence interval. Diagonal graphs show the kernel density estimate of each parameter, to illustrate skewness and multimodality in the data. Dots in the top right triangle represents the correlation between two parameters (blue for negative correlations, red for positive correlations) displaying the Pearson correlation coefficients, as shown in details on Fig. S8. As analgesia, phosphorylation and respiration were measured in independent cohorts, the correlation is computed at the cohort level. As respiration was measured in the first 30 minutes (Fig. S8B), correlations are shown in this time window. There are two anticorrelated clusters of quantities (respiratory depression, mobility, CBV vs. FC index, phosphorylation, analgesia). (**C**) Average physiological readouts from the morphine 70 mg/kg session shown as a z-score heatmap for all sub-phases (**left**) and corresponding correlation matrix (**right**). (**D**) Loadings (**left**) and explained variances (**right**) from principal components of the correlation matrix shown in (**C**). We retained principal components whose explained variances were above those predicted from random white noise. (**E**) Projection of the average physiological readouts from all sessions on the two principal components retained from (**D**). The first principal component (PC1) indicates increased FC index, analgesia and phosphorylation. Notably, sessions are sorted along PC1 by increasing dose, with baseline sub-phases (BP 1/BP 2) and all saline sub-phases are tightly grouped with highly negative loadings on PC1. The second principal component (PC2) indicates increased CBV and mobility. Sub-phases directly following the injection of morphine are tightly grouped with highly positive loadings on PC2.

Importantly, these vascular and behavioral events showed poor temporal correlation with the changes in functional connectivity (Fig. S8), which was delayed by about 20 minutes at the 30 mg/kg dose (Fig. 6A). In addition, similar connectivity changes were observed in BALB/cJ mice further demonstrating that changes in locomotion, perfusion and FC are predominantly independent effects of opioids (Fig. S9). Thus, while the kinetics of locomotion, respiration and regional CBV changes are fast and synchronous, morphine-induced reorganization of FC is relatively slow, as compared to these three parameters. Morphine-induced analgesia, a therapeutically important parameter measured in the hot plate test in a parallel awake mice cohort, displayed kinetics similar to the relatively slow FC changes (Fig. 6A and B and Fig. S7A). Additionally, these relatively slow changes in FC patterns and analgesia were also well-correlated with changes in the activation level of MOP at the whole brain scale, as detected by the proxy measurement of morphine-specific phosphorylation of MOP at the S375 residue (Fig. 6A and B and Fig. S7D), measured in another parallel awake mice cohort. These data suggest the existence of two latent processes driving independently rCBV, respiration and locomotion in a fast and transient manner while FC changes, analgesia and MOP activation appear later in time. To confirm these observations, we performed a principal component analysis of all physiological readouts but respiratory depression from session morphine 70 mg/kg (Fig. 6C – E). We found that two principal components are sufficient to explain most of the readouts variance. The first component is most correlated with the FC index, analgesia and phosphorylation, while the second is most correlated with motion and CBV (Fig. 6D). Finally, we projected the physiological readouts from the saline and morphine sessions onto the two previously computed principal components. As expected, the first principal component (PC1) sorts sub-phases by increasing dose, with sub-phases 4 to 6 having higher loading along PC1 than earlier sub-phases. Notably, all baseline sub-phases from morphine sessions and all saline sub-phases are tightly grouped with highly negative loadings along PC1. Conversely, sub-phases directly following morphine injections have highly positive loadings along the second principal component (PC2).

Taken together, our results show that major opioid drugs lead to significant changes in region-wide FC patterns, through the modification of correlative behavior in large neuronal assemblies (Fig. S8). These FC alterations are pharmacologically relevant since they are MOP-specific and encompass timing, dose-dependence, and the development of tolerance due to prolonged drug exposure. Importantly, these FC alterations, but not local changes in perfusion, are well-correlated with the development of analgesia, a therapeutically important behavioral read-out, but not with respiratory depression, a serious side-effect of opioids.

## DISCUSSION

In this study, we have developed a multimodal and minimally-invasive approach to assess drug-induced functional alterations in the awake rodent brain with high sensitivity, through the intact skull. Our findings reveal that major opioids lead to a remarkably consistent dysconnectivity pattern, referred to as the “opioid fingerprint”. This pattern predominantly involves the emergence of anticorrelations between cortical and subcortical regions, resulting from altered oscillatory patterns, gradually occurring following opioid administration. Importantly, these dysconnectivity patterns are strongly correlated with and may potentially underlie the development of analgesia, which represents the primary therapeutic effect of opioids.

### Methodology

In this pioneering study our primary focus was to ensure robustness and reproducibility, prompting specific technological trade-offs that can be readily addressed in future investigations. One key trade-off encountered in awake animal fUS imaging pertains to the balance between invasiveness and sensitivity. Employing intact skull imaging results in a modest overall loss of the power Doppler signal, which becomes more pronounced in regions located under skull sutures. An alternative approach involves craniotomy, potentially accompanied by a plastic prosthesis (*11, 24*) to counteract the loss of sensitivity. However, this entails more invasive surgical procedures, modifies cortical blood flow patterns, and introduces increased variability due to post-operative complications. An intermediate solution is skull thinning (*10, 16, 25, 26*), which is less invasive but may still induce post-surgical variability at the cohort level. Our comparison with the literature suggested that transcranial fUS imaging, in our experimental setup, remains at least as sensitive as state of the art preclinical fMRI, making it well-suited for an initial assessment of drug-induced effects in the awake mouse brain with low variability.

The second trade-off, partly interconnected with the first, involves the compromise between spatial coverage, temporal resolution and experimental throughput. To validate pharmacological effects across various mouse strains and several drugs, including two of them at different doses, we opted to image a single representative brain slice, easily identified at the beginning of the imaging sessions based on its specific vascular landmarks. Remarkably, this single slice, sampled at up to 2 Hz, yielded already highly significant and reproducible outcomes. Our imaging protocol can be readily extended further for 3D coverage employing 2D matrix or RCA probes (*14, 27, 28*). However, our unpublished results indicate that matrix probes still require development to obtain the requisite sensitivity for minimally-invasive transcranial imaging of pharmacological responses in mice. Currently, single slice 2D imaging with a linear array probe provides the highest temporal resolution for fUS imaging, motivating our experimental choice. Importantly, recent results indicate that 2D sections of 3D resting-state networks (RSNs) coincide well with RSNs obtained from 2D power Doppler images (*29*), suggesting that spurious latent network correlation structures reported in under sampling methods, such as ours, represent a powerful and economical approach to detect dysconnections at the whole-brain level.

### Opioid-induced dysconnectivity

The functional connectivity changes detected in our sample coronal slice report changes in correlations of the spontaneous fluctuation of blood flow. On their own, these results do not allow to draw conclusions about the precise mechanism causing this loss of synchronicity. Changes in neural dynamics (local shift of E/I or changes in synaptic weights), out of plane connectivity changes, as well as non-neuronal sources, like neuro-vascular decoupling could separately or together explain the change in the reported functional connectivity patterns. Independently from their exact origin, our results suggest large-scale reorganization of functional connectivity patterns, which are relatively slow (at around 20 min for morphine), following the administration of opioids. While the predominant effect is a decrease in the zero-lag Pearson’s correlation of the Power Doppler signal between the thalamus and cortical regions, accompanied by an important change in hippocampal FC patterns, dynamical analysis suggests that this dysconnectivity is not a simple loss of information exchange between cortical and subcortical regions, but rather a reconfiguration of coupled oscillatory patterns. Indeed, morphine-induced dysconnectivity displays a rich spatio-temporal pattern, but notably, highly-significant and reproducible responses can already be detected by focusing solely on the classical zero-lag correlation analysis.

At steady-state, integration of a large spectrum of oscillatory patterns leads to the characteristic power-law spectrum or 1/f statistics in various experimental read-outs (obtained for example by electrophysiology or EEG), suggesting self-organized criticality of brain dynamics (*20*). This may enable rapid and robust large-scale reconfiguration of functional networks in response to exogenous stimuli. Notably, the power-law can be observed in the spectrum of the power Doppler signal of our control cohorts (saline treatment). Perturbations, such as sensory stimuli or motor output, were proposed to reduce the critical state in the cortex and provide transient stability through oscillations (*19*). The resulting rhythmic cortical feedback to the thalamus is a major factor in the amplification of thalamocortical oscillations and with increasing commitment to an oscillatory network, the responsiveness of neurons to external inputs progressively decreases (*19*).

Our results indicate that pharmacological treatment may induce similar critical-state-reducing perturbations, since morphine treatment results in important changes both in the frequency repertoire and in the temporal organization (i. e. lag structure) of thalamo-cortical networks. Thus, by perturbing cortical critical-state dynamics through modification of the excitation/inhibition balance (*30*), MOP acting on inhibitory interneurons (*31*) may induce specific oscillations to reduce the neuronal responses to prominent environmental influences, such as pain. As slow, large-scale oscillations are considered as modulators of faster local events, the emergence of the slow morphine-induced oscillatory activity at 0.05Hz may also alter the power of faster gamma oscillations, which are reportedly modulated by morphine in hippocampal slices (*32*) or in the awake mouse striatum (*33*). Future research combining electrophysiology and fUS imaging techniques in awake mice (*34*) may disentangle the intricate relationship between opioid-modulated local and large-scale events and their implication in pain perception and analgesia.

To the best of our knowledge, only a single study has thus far investigated the acute effect of MOP agonist on functional connectivity in mouse models (*8*) by measuring the influence of oxycodone (2 mg/kg) on BOLD activation and zero-lag FC patterns in mice under dexmedetomidine anesthesia. Through independent component analysis (ICA), they identified 71 components constituting a connectivity network encompassing the entire brain. Notably, their findings primarily revealed an overall reduction in FC. Furthermore, they show region-specific increased perfusion, which correlates with MOP-enriched clusters (periaqueductal gray, substantia nigra, caudate putamen and nucleus accumbens). Both effects are absent in MOP knockout mice. Notably, in contrast to our results, this study does not report opioid-induced changes in cortical-subcortical FC, which is known to be affected by anesthesia (*35*). The validity of our findings is reinforced by similar observations in an awake human fMRI study, where morphine- and alcohol-induced connectome changes were compared to placebo controls (*36*). Utilizing an advanced model based on 8 canonical resting-state networks, the most notable finding in their investigation is a dysconnectivity or induced anticorrelation between the somatosensory cortex and the hippocampus, which is consistent with our own findings. Additionally, fMRI imaging of heroin-maintained outpatients demonstrated that acute injection of heroin, a pro-drug of morphine, induces cortico-thalamic disconnection (*37*), and multiple studies reported opioid-induced dysconnectivity between the cortical and subcortical regions (*38, 39*). In conclusion, our fUS imaging-based reporting of cortico-subcortical dysconnection as the primary effect of morphine likely represents a significant and clinically relevant outcome. Moving forward, as we extend mouse opioid fingerprinting to the whole mouse brain, we anticipate the potential for new findings that could be translated from mouse models to humans (*40*), owing to the functional similarities between BOLD and fUS imaging. Such cross-species insights hold promise for advancing our understanding of opioid-related effects and their implications for human neuroscience and clinical applications.

### Opioid-induced CBV changes

Our results also demonstrate MOP-induced local CBV changes in several regions following opioid injection. This effect exhibits a bi-phasic pattern, characterized by an initial hyperperfusion followed by hypoperfusion, with kinetics showing significant dependence on the type and dose of the agonist. Previous pharmaco-fMRI studies have reported changes in the BOLD signal following acute injections of oxycodone in awake restrained mice (*41*) and in awake rats (*42*) as well as buprenorphine in awake rats (*43*). The activation and deactivation patterns reported in these studies exhibit considerable variability, with a focus on the hyperacute effects of the compounds occurring within a few minutes after injection. Given the dose-dependence and bi-phasic nature of the response reported in our present study, we emphasize the importance of having not only precise time resolution but also a prolonged longitudinal follow-up of hemodynamic changes. Indeed, a clinical study that longitudinally tracked BOLD intensity changes following injection of a fentanyl analog revealed region-specific local activations with rapid kinetics lasting only a few minutes (*44*), which aligns with our own results. However, our technical advancement, employing fUS imaging and awake preparation, allowed us to demonstrate that these transient perfusion changes are strongly correlated with two behavioral read-outs, hypermobility and respiratory depression, but not with the clinically critical read-out of analgesia, which spans longer time scales. This difference in dynamics and the relatively late changes in functional MOP effects are particularly intriguing in the light of the functional selectivity hypothesis (*27*). Overall, these results suggest that opioids can selectively activate specific effector cell populations and/or signaling pathways through the MOP, leading to distinct downstream effects, which allows to clearly distinguish analgesia and respiratory depression through their differential correlation with CBV and FC changes. Future research utilizing fUS imaging in mouse strains with targeted inactivation of individual components of the MOP signaling pathway will enable functional dissection of these effects, providing deeper insights into the mechanisms underlying opioid-induced alterations in brain hemodynamics and behavior and a putative future path to develop efficient analgesics with reduced side-effects.

### Conclusion

In conclusion, we present a multimodal experimental approach that allows the quantitative tracking of drug-induced functional “fingerprints” of brain activation and functional connectivity, in correlation with behavior, in the awake and behaving mouse. Our findings highlight the dysconnectivity between cortical and subcortical regions, temporally correlated with MOP activation and the development of analgesia, as the prominent feature of the MOP-specific fingerprint.

Our approach has the potential to contribute to bridging the explanatory gap between cellular/microscale neuronal mechanisms and large-scale neural dynamics that underlie behavior. By understanding the correlations between molecular mechanisms, behavior and functional signatures controlling opioid-related effects and side-effects, we can gain valuable insights into alternative therapies for pain management, tolerance, addiction/dependence, withdrawal and respiratory depression. In the future, this approach holds promise in facilitating better prediction of the effects and therapeutic windows of newly developed drugs, while aiding in the selection of appropriate application areas based on the balance between analgesia and acceptable side effects. Ultimately, our research may contribute to the advancement of opioid-related therapeutic strategies to improve patient outcomes.

## Material and Methods

### Animals

Experiments were performed on JAX^TM^ C57BL/6J mice obtained from Charles River Laboratories (DE) or from Janvier Labs (FRA). BALB/cJ mice were obtained from Janvier Labs (FRA). MOP knockout (MOP^-/-^) mice were original provided by Dr. H. Loh (University of Minnesota, Minneapolis, MN), breaded and backcrossed to WT control JAX^TM^ C57BL/6J mice obtained from Charles River Laboratories (DE) for more than 12 years in the Institute of Pharmacology and Toxicology at the University Hospital in Jena (DE). Knock-in mice expressing an HA-epitope tag MOP (HA-MOP, Oprm1^em1Shlz^, MGI:6117675) were provided by Stefan Schulz (University Hospital Jena, Institute of Pharmacology and Toxicology, Jena, DE) and described in detail (*45*).

Animals were housed 2 – 5 per cage under a 12–hour light–dark cycle with ad libitum access to food and water. In all experiments, we used cohorts of n = 10-12 male mice aged 7 – 14 weeks and weighing 18 – 24 g at the beginning of the experiments. Experimental procedures were performed in accordance with the Council of European Union directive of September 22, 2010 (2010/63/UE) and with the French decree of February 1, 2013 (n°2013-118) and were approved by Thuringian state authorities and by the French *Comité d’éthique en matière d’expérimentation animale* (APAFIS#25912-2019072317104707 v2). Our study is reported in accordance with the ARRIVE (Animal Research: Reporting In Vivo Experiments) guidelines (*46*).

### Drugs and routes of administration

All drugs were freshly prepared prior to use and were injected subcutaneously at a volume of 10 µl/g body weight. Drugs were diluted in 0.9% (W/V) saline for injections. Drugs were obtained and used as follows: morphine sulphate (10 – 70 mg/kg; Hameln Inc., Hameln, Germany), fentanyl citrate (0.1 – 0.3 mg/kg ; Rotexmedica, Trittau, Germany), levomethadone hydrochloride (10 mg/kg; Sanofi-Aventis, Frankfurt, Germany), buprenorphine hydrochloride (3 mg/kg; Indivior, Mannheim, Germany) and naloxone hydrochloride (2 mg/kg; Ratiopharm, Ulm, Germany). For chronic infusion with Alzet osmotic minipumps (1007D), fentanyl citrate salt (2 mg/kg/day) and morphine sulphate salt pentahydrate (17 mg/kg/day) were obtained from Sigma-Aldrich (St. Louis, MO). Drugs were diluted in phosphate-buffered saline for acute injections or dissolved in sterile water and then diluted in phosphate-buffered saline for osmotic pump delivery.

### Behavior studies

For consistency, one experimenter and a dedicated assistant performed all *in vivo* drug administrations and behavioral testing. All testing was conducted between 7am and 4pm in an isolated, temperature- and light-controlled room. Mice were acclimated for at least 2 weeks before testing. Only the experimenter and assistant had free access to the room and entered the room 30 min before commencement of testing to eliminate potential olfactory-induced changes in nociception. Animals were assigned to groups randomly before testing. Mice were excluded from the study if they displayed any bodily injuries from aggressions with cage mates. The experimenter was blinded to treatment and/or genotype throughout the course of behavioral testing. All drugs were given to the experimenter in coded vials and decoded after completion of the experiment.

### Tolerance paradigms

Acute effects of 70 mg/kg morphine or 0.3 mg/kg fentanyl were recorded. To induce opioid tolerance, mice were implanted with Alzet osmotic minipumps containing the same drug that was used in the acute dose. Osmotic minipumps delivered total daily doses 17 mg/kg morphine or of 2 mg/kg fentanyl at a rate of 0.5 µl/h. On day 7, mice were again treated using a repeated acute dose with morphine or fentanyl.

### Osmotic pump implantation

A single 1007D Alzet osmotic minipump (Charles River Laboratories, DE) was subcutaneously implanted on the left limb of each mouse under light isoflurane anesthesia and meloxicam analgesia. A small incision was made in the skin on the mouse’s left flank with a scalpel, a small pocket was formed just beneath the skin, and the minipump was inserted. The incision was closed using 7 mm wound clips (Charles River Laboratories).

### Surgical procedure and preparation for imaging

The mice were prepared following a previously published protocol (*16*). The procedure consists in the surgery for the implantation of a metallic headplate (Neurotar, Finland) and the habituation protocol to head fixed condition before imaging sessions.

### Headplate surgery

The mouse was pre-treated with Meloxicam (4 mg/kg) before surgical ketamine/xylazine (100 mg/10 mg/kg) anesthesia. After the depth of anesthesia was verified the mouse was positioned onto the stereotaxic frame. The body temperature at 37°C was maintained by a heat blanket during the complete duration of the procedure. Lidocaine (2%, 0.2 mL) was administered subcutaneously under the scalp skin before incising the skin. After removing the skin from behind the occipital bone to the beginning of the nasal bone, to obtain an imaging window of 13 mm x 21 mm, the skull was carefully cleaned with iodine solution to remove the remaining periosteum. The temporalis muscles were detached to obtain the widest imaging window possible. By using the headplate as a template, two superficial holes (1 mm diameter) were drilled in the front and the back of the skull to position the anchoring screws. A specific care is given to avoid drilling through the skull. Afterwards the headplate was screwed to the skull and the screws fixed with fast solidifying surgical cement (Superbond C&B) to firmly attach the implant. Finally, the skull was protected by a thin layer of surgical glue (Vetbond, 3M, USA) to protect the bone and to close the wounds surrounding the imaging window. The animal was removed from the stereotaxic frame after the cement dried and left to recover in an individual heated chamber.

### Handling and Habituation

After surgery the animal was left for 4 to 6 days to recover and then was daily handled and trained to get used to head fixed condition in the Mobile HomeCage® (MHC) as described as follows. During the first 2 days the animal is handled in the experimental room for habituation to the environment and the experimenter. On the 3^rd^ day the animal is left 5 – 10 min to freely explore the MHC. After the fourth day the animal is clamped daily increasing the duration of head fixed condition of 5 min per day until reaching 30 min. On the 7^th^ day some gel is poured on the imaging window without imaging. On the 8^th^ day the probe is lowered down. On the last day the animal is imaged to obtain an anatomical reference of its brain vascular tree.

### Pharmaco-fUS imaging protocol

The mouse was weighted and clamped to the MHC apparatus. The imaging window was humidified with saline and ultrasound gel was applied on top of the skull. After the probe was lowered down until it contacted the ultrasound gel, the machine was turned on in live view for positioning the probe above the reference plane based on the anatomical vascular landmarks. In general, recordings lasted 20 min for baselines followed by multiple 30 min recordings (typically 2 for 1 h follow-up). After each acquisition period, data was transferred to avoid saturating memory, leaving 5 min of dead time between two recordings. Injections were performed between two consecutive recordings to avoid injection-induced movement artifacts during the recording. The animal was injected subcutaneously while being head-fixed. Tracking was recorded using MHC Tracker software (Neurotar, Finland).

### fUS Imaging

A 15 MHz ultrasound probe (15 MHz, 128 elements, 110 µm pitch) connected to a prototype functional ultrasound scanner (Iconeus, Paris, France and Inserm ART Biomedical ultrasound) was used. The positioning of the probe above the head of the animal was performed using a motorized probe holder allowing translations along three axes (1µm precision) and rotation around the vertical axis (0.1° precision). A previously described acquisition sequence was used; 11 tilted plane waves ranging from -10° to 10° by increments of 2° were fired at a pulse repetition frequency (PRF) of 5500 Hz, leading to a 500 Hz sampling rate after beamforming and compounding (*15*). Power Doppler images were then computed offline using ensembles of 200 compound frames, following manual skull-stripping and SVD-based clutter filtering (reconstruction of the ensembles frames by zeroing out the first 60 singular values) (*47*). A delay of 0.1s is added between each ensemble to reduce the risk of memory saturation during long recordings. The resulting power Doppler acquisitions are sampled at 2 Hz.

### Power Doppler Template

After data format standardization (see Supplementary Methods), we computed a template for each imaging plane following the Allen Brain Institute methodology (*18*). All images were rigidly registered using Elastix (*48*) to a single reference image of the dataset. Images were split in left and right hemispheres and each hemisphere was independently mirrored to obtain two datasets of left and right hemispheres. Right hemisphere images were then affine (translation+rotation+scaling+shearing) registered on left hemisphere images and an average of all images was computed to obtain the first iteration of a template. All images were again affine registered to the first template iteration and averaged to produce a second iteration. Finally, all images were non-affine registered to the second template iteration and averaged, leading to the third and final template image. This final template was used for co-registering all power Doppler acquisitions. Rigid and non-rigid transform parameters were systematically investigated in a grid search to find optimal values (see Supplementary Methods).

### Power Doppler Preprocessing

Power Doppler acquisitions were preprocessed in Python using Nilearn 0.10.1 (*49*) to mitigate motion artifacts. A scrubbing strategy was first applied to reject frames displaying strong motion artifacts (*50*). Frame rejection was based on the DVARS metric (*50*), computed on the first five temporal singular vectors obtained via singular value decomposition (SVD) of the centered (time, voxels) power Doppler matrix. DVARS was computed on these singular vectors and not on the whole data matrix as they better represent sources of global variations likely due to motion artifacts. The resulting metric was used to generate a sample mask by thresholding it at 0.5 standard deviation. The sample mask was then convolved with a kernel of size 2 to remove samples directly following each rejection period. Samples outside of the sample mask were then replaced by the temporal-average computed from the data inside of the sample mask. This “pre-scrubbing” step is performed to remove motion artifacts likely to produce ringing artifacts in the subsequent temporal filtering step, while keeping a stable sampling frequency. Notably, higher-order interpolation methods routinely used in fMRI "pre-scrubbing" can be detrimental considering the high proportion of motion artifacts in awake mice data. The data was then standardized to z-scores and the first temporal singular vector was computed and included as a confound signal. The first temporal singular vector was used instead of the global signal as it better represents sources of global variations. Confound regression and temporal filtering ([0.01, 0.1] Hz) was then performed. Finally, the sample mask was used to reject frames that were previously replaced by the mean.

### Automatic parcellation

In order to focus on FC changes in regions that show the highest level of FC at the baseline, an automatic parcellation approach has been used. A voxel-wise correlation matrix was computed, by using a seed computed from a square of 9 voxels surrounding the voxel of interest. The matrix rows were then clustered using the K-means algorithm from scikit-learn (*51*) with n = 18 clusters. Clusters intersecting with the midline or the border of the image were considered artifacts and rejected. For coronal slices, the parcellation was symmetrized along the midline. Each ROI was deflated by 1 voxel to reduce the influence of potential voxel mis-registration from one ROI to another. Details on the procedure and its validation are given in the (Fig. S2).

### Correlation matrices and seed-based maps

Cleaned time series from each ROI were averaged and correlated (Pearson correlation) over 10-minute windows to produce correlation matrices (*15*). Seed-based maps were also computed by voxel-wise correlation with each ROI average time series. Both correlation matrices and seed-based maps follow a mass univariate approach. Correlation matrices and seed-based maps from each experimental cohort were compared to a reference cohort (n = 47) of animals injected with saline and imaged in the same conditions. Comparisons were made for each 10 minutes window. Samples of correlation coefficients were compared using the Mann-Whitney U-test (*52*) following Fisher z-transformation. The resulting p-values were corrected for multiple comparison using the Benjamini-Hochberg step-up procedure (*53*) with a FDR threshold set at 0.05.

### Circular graphs

*Circular graphs* are alternative representations of correlation matrices and are intended for better visualization of FC changes. Networks were plotted using the NetworkX library (*54*).

In average graphs, each node represents a ROI and each edge represents the averaged Pearson correlation coefficient between two nodes. The value for both nodes and edges is represented as increased opacity in red for positive correlations and blue for anti-correlations. For better readability, weak edges for correlation whose absolute value is below 0.3 are not displayed. The node color encodes for the regional global connectivity (rGC) (*55*), *i.e.* the sum of the correlation between one node and all other nodes. The rGC is normalized relatively to the ROI-wise maximum average rGC during the reference condition after saline injection. This normalization highlights differences in rGC.

In significance graphs, edges represent the difference of correlation between the experimental condition and the reference cohort. Significance graphs only display edges whose differences are significant (see previous paragraph). Nodes represent the sum of the connectivity difference independently of the effect significance. The node values are normalized to highlight difference of the effect by half of the maximum (ROI wise) average GC during reference condition after saline injection.

### FC index

The functional connectivity index was established by using the maximal morphine dose (70 mg/kg) as a reference (Fig. S10). Correlation matrices are estimated for each animal and each 10-minute window. Singular value Decomposition (SVD) is then performed on the average matrices over the full cohort. Singular value decomposition is performed on independent edges (inferior left triangle of the average matrix) across windows (n_edges_ x n_phases_ = 153 x 8). The second component of the non centered SVD reliably represents the morphine effect (Fig. S9A and B). We calculate the FC index by projecting the individual correlation matrices on this second (in terms of ROIs pairs or matrix edges) component. The index unit is expressed as a Pearson’s correlation coefficient and can be compared across experimental conditions. The FC index dynamically represents the intensity of the morphine fingerprint for each individual animal.

### Tracking

During awake fUS imaging acquisitions, animal movements are tracked using the Neurotar Mobile HomeCage® tracking system (*56*). The system estimates the animal position through the movements of the cage at a sampling rate of 100Hz and a precision <1mm. Mobility measures are resampled to the power Doppler sampling rate and the instant speed is computed as the absolute value of first order derivative of the resampled position. Based on a meta-analysis of 186 recordings at baseline, we defined the threshold of movement at the elbow of the distribution of speed (Fig. S12). Thus, the animal is considered as moving when its speed exceeds 5 mm/s.

### CBV

The power Doppler (PD) signal is proportional to CBV. To represent the evolution of perfusion at the cohort level, recordings from the same session are concatenated and each voxel value is normalized based on the last 2min of the baseline which results in ΔCBV = (PD-<PD>)/<PD> where <PD> is the average value over the last 2min of baseline. ΔCBV is represented at the regional level by averaging all voxels of the ROI. The cohort level perfusion is represented as the average perfusion across all individuals for each timestamp in the form of a carpet plot whose saturation is [-0.5, 0.5].

### Lag analysis

Lag analysis consists in computing seed-based maps using time-delayed versions of a given seed. When the delay is positive, the seed time series is shifted forward in time (Fig 3A, left). In this case, a high correlation coefficient between a voxel and the positively delayed seed could indicate that events in the seed time series precede those in the voxel time series. When the delay is negative, the seed time series is shifted backward in time (Fig 3A, right). In this case, a high correlation coefficient between a voxel and the negatively delayed seed could indicate that events in the voxel time series precede those in the seed time series. Because coherence of the correlation structure decays with increasing lag, the maps at each lag are scaled (divided by the standard deviation) in order to be comparable in terms of distribution. Second order statistics are computed similarly as described above for SBMs on the cross-correlation value before normalization.

### Spectrum analysis

For the analysis of CBV fluctuation frequency spectra, signals are minimally processed: the raw Power Doppler is expressed as percent signal change (ΔCBV), relative to the mean value. Then regional average haemodynamic is extracted for each animal and the power spectral density is computed via Welch’s method, using the scipy.signal.welch from the SciPy Python package (*57*). The average normalized spectrum is displayed with 95% confidence intervals in log-log diagram. The slope of the 1/f signal is computed using the numpy.polyfit from the Numpy Python package^56^ on the FC frequency band [0.01, 0.1] Hz.

### Data Selection

Our experiments were performed on cohorts of n = 10 mice for C57BL/6J, n = 11 mice for MOP^-/-^ and n = 12 mice for BALB/cJ. Recordings were discarded from further analysis because of corrupted files (n = 3/296), animal death (n = 1/296) and mis-registration (n = 8/296). In addition, aberrant FC values inducing significant differences from our saline cohort at baseline resulted in the rejection of two recordings in the naloxone experiment. This baseline effect could be associated with extreme mobility values (>=50% against 20% on average). A summary table in the supplementary methods recapitulates the analyzed data for fUS experiments (Tab. S1).

## Supporting information

Suppplementary Materials and Methods

## Acknowledgements

We thank Monique Brendel, Madlen Hüttl, Barbara Herböck, Hippolyte de Valmont and Adam Cellier for excellent technical assistance. We acknowledge the ElfUS platform of IPNP directed by Luis Villanueva and Laurence Bourgeais Rambur during this project as well as the work of the animal facility staff of IPNP under the supervision of Tifanny Leclerc for the care of the animals. We thank Gyorgy Buzsaki for discussion of the dynamical fUS data and Diana Zala for discussion of the manuscript. We thank Bruno Osmanski and Luc Eglin from Iconeus for support and discussion on analysis.

## Funding

Else Kröner Fresenius Stiftung (2019_A68) to A.K

Interdisciplinary Center for Clinical Research Jena (AMSP 03) to A.K.

EMBO Short-Term Fellowship (n° 8439) to A.K.

PhD scholarship from Boehringer Ingelheim to J.C.M.

DAAD short-term research grant (57507442) to J.C.M.

Inserm Biomedical Ultrasound ART project to Z.L.

## Author contributions

A.K., J.C.M. and L.B. contributed to the surgery, habituation and fUS imaging. A.K. performed and analyzed all *in vivo* behavioral pharmacology studies. J.C.M. and S.D. created and implemented a custom functional connectivity analysis pipeline. J.C.M. and S.D. analyzed all imaged data. RS contributed to implementation of the study and discussed the results. S.S. provided the animals and contributed to the scientific discussion. M.T. and T.D developed and provided the imaging setup and acquisition software, provided technical and scientific assistance on acquisition and analysis and contributed to the discussion. The manuscript was written and revised by Z.L., A.K., J.C.M., and S.D., with editing and suggestions from the other authors.

## Competing Interests

T.D., M. T and Z. L. are founders and stakeholders of Iconeus. S.D. is an employee of Iconeus. S.S. is the founder and scientific advisor of 7TM Antibodies GmbH, Jena, Germany. All other authors declare no competing interests.

### Author Information’s

Readers are welcome to comment on the online version of the paper. Correspondence should be addressed to A.K. and Z.L.

## Data and code availability

The pre-processed power Doppler data that support the findings of this study are available in Zenodo with the identifier … *(The Zenodo DOI will be active at publication and full data access is available to editors and referees upon request.)*

The source codes used for the generation of our figures and statistical analysis are available in Zenodo with the identifier … *(The Zenodo DOI will be active at publication and full code access is available to editors and referees upon request.)*

## Supplementary Materials

Materials and Methods

Supplementary Text

Figs. S1 to S12

Tables S1

Video S1

