## Supplementary material for "Opioid-Induced Inter-regional Dysconnectivity Correlates with Analgesia in Awake Mouse Brains": Suppplementary Materials and Methods

### Cortico-Subcortical Dysconnectivity Following Opioid Administration Correlates with Analgesia in the Awake Mouse Brain

and

Zsolt Lenkei, Institute of Psychiatry and Neurosciences of Paris, INSERM U1266, Université Paris-Cité, 102-108 rue de la Santé, F-75014 Paris, France, Phone: +33-1-40789251,

#### EXTENDED DATA METHODS

##### *Templates and regions of interest*

Statistical analyses used in the current study rely on voxel-wise comparisons, requiring properly aligned brain volumes. Recent advancements in powerful registration methods for power Doppler images have shown promising potential[1,2]. These tools are currently being evaluated for their effectiveness in transforming power Doppler images to the CCFv3 coordinate system (Allen space)[3]. Leveraging the high redundancy and reproducibility of our datasets, we opted to use a custom tailored-made pipeline for brain positioning, utilizing a data-driven approach. Our pipeline contains three steps:

1. *Reproducible positioning of the probe.* We identified specific landmarks of the vascular tree, allowing reproducible probe positioning when used as references during imaging sessions, even across experimenters.
2. *Post-hoc registration*. We generated a symmetrical power Doppler template of our plane of interest based on 526 images of the same plane acquired across 206 individual mice.
1. *Data-driven automated parcellation.* In order to maximize the functional relevance of the investigated connectivity networks, we generated ROIs by voxel-wise clustering of the baseline functional connectivity matrix.

##### *Positioning*

For 2D imaging, the probe placement is critical, as no registration can rescue misplaced data. This positioning is facilitated in our case by the head fixation. As the implanted metal headplate is parallel to the cortical surface all brains share a common space, with an 8° angle in the coronal plane with respect to the Paxinos mouse brain atlas (ref. Paxinos et al.) and an 18° angle with respect to the Allen mouse brain atlas[3]. The standard acquisition plane is obtained by manual adjustment, through live visualization of the branching point between the internal carotid and the anterior choroidal arteries (Extended Data Fig. 1). This branching point is easily identified by the experimenter with a precision of ~100µm, providing a reproducible antero-posterior vascular reference point. Other anteriorities may then be obtained with respect to this reference.

##### *Post-hoc registration to a custom template*

526 images were used to generate a single reference template of our plane of interest. These sessions were recorded over 206 individual mice, with several mice imaged multiple times. Below we detail the different steps of the procedure briefly described in Methods (Power Doppler Template):

1. For each of the dataset session:
    a) All the images in the session are concatenated along the temporal axis $\{I_i, i \in [1, N]\}$
    b) The average is exported as a simple 2D image. $\{\hat{I}_i, i \in [1, N]\}$
2. The «best image» is selected as the first reference (Extended Data Fig. 1).
3. All images are registered to the first reference using an affine transformation. $\{\hat{I}_{I0,i}, i \in [1, N]\}$
4. All affine registered images are split in half along the vertical axis and both halves mirrored to create symmetrical images of each halve.
5. (Left-left, Right-Right). $\{(\hat{I}_{I0,i,L-L}, \hat{I}_{I0,i,R-R}), i \in [1, N]\}$
6. All Right-Right images are registered to their Left-Left counterparts using an affine transformation.
7. $\{(\hat{I}_{I0,i,L-L}, (\hat{I}_{I0,i,R-R})_{\hat{I}I0,i,L-L}), i \in [1, N]\}$
8. A second template is generated by averaging all symmetrical images.. T2
9. All images are then registered to T2 using a b-spline transformation. $\{((\hat{I}_{I0,i,L-L})_{T1}, ((\hat{I}_{I0,i,R-R})_{\hat{I}I0,i,L-L})_{T1}), i \in [1, N]\}$
1. The final template T3 is obtained by averaging these transformed images.

***Data-driven automated parcellation***

For the definition of ROIs, a data driven approach was performed, as briefly described in Methods (Automatic parcellation, Extended Data Fig. 2). In details, functional ROIs were defined as voxel clusters showing high functional connectivity at baseline. By virtually scanning the image with a small seed, composed of the seed voxel and immediate neighbouring voxels (9 voxels in total), a voxel-wise correlation matrix was build, followed by k-means clustering to identify regions of common functional connectivity territories. By gradually elevating the number of clusters, we found that on our plan of interest more than 21 clusters lead to the emergence of ROIs without bi-lateral symmetry. Clusters touching the edges of the image, including the midline low signal-to-noise zone, were discarded. The resulting, mostly symmetric ROI map was further symmetrized to ensure proper localization of bilateral voxels after registration on a symmetric template. Finally the ROIs were deflated to account for small misregistrations, resulting in the final number of 18 conserved individual ROIs.

In order to evaluate the validity of our parcellation, we have verified how cluster edges correspond to intensity changes (i.e. borders) in the voxelwise global connectivity map (sum of the absolute value of the correlation coefficient of one voxel with all the other voxels), computed as:

$$GG_{i,j} = \sum_{[k,l]\neq[i,j]} | < s_{i,j}, s_{k,l} > |$$

Importantly, main anatomical regions, such as cerebral cortex, hippocampus, dorsal thalamus, vetral thalamus and hypothalamus) can be identified in the global connectivity intensity distribution. Laplacian filtering (2D isotropic measure of the 2nd spatial derivative of an image, used for edge detection) of the global connectivity map show distinct borders of connectivity distributions Extended Data Fig. 2c, II.) and importantly, the ROIs selected above fit within this borders (Extended Data Fig. 2c, III.), but not fit within high-intensity regions in the Power Doppler image, showing that ROIs correspond well to the functionally most connected regions. Finally, superposition of our data-based ROIs on the Allen Mouse Brain Atlas shows excellent anatomical correspondence both in global and local organization, as ROIs do not overlap neighboring regions such as the cortex and hippocampus and even delineate functional units inside these regions such a the somatosensory cortex and visual association area, the hippocampal CA1 and CA3 regions, the lateral posterior nucleus and posterior complex in the polymodal Thalamus and the lateral geniculate nucleus in the unimodal thalamus.

**Tracking**

Tracking data was acquired using the Neurotar embedded tracking system. By detecting the position of two magnets attached to the floating cage with high precision (100Hz, <1mm), a cinematic model calculates the virtual head position of the mouse (Extended Data Fig. 10, left). Because of this high frequency, instantaneous speed is subject to high frequency instability of the derivative. To mitigate the high-frequency induced instability of the instantaneous speed derivative, we re-sampled cage positions to the fUS signal timestamps (2Hz), and virtual animal velocity was re-estimated using these down-sampled time series. The resulting velocity estimate is more stable during low movement epochs. We computed over 186 recordings of the the baseline cumulative distribution of mobility and, by using the Kneedle algorithm[4] for elbow detection, we identified a regime transition, putatively useful as a threshold of mobility detection (i.e. the ratio of time spent above a certain speed) at around 3.86mm/s. We adopted a conservative threshold of 5mm/s for mobility estimation.

**Functional connectivity index**

In order to construct a functional connectivity (FC) index, i. e. a single value representative of the morphine effect on functional connectivity to calculate FC correlations with other read-outs such as CBV or behaviour, we used multivariate analysis, by performing singular value decomposition (SVD) on the average FC matrix after morphine injection (70 mg/kg). $X = \{FC_{i,j}, \forall i, j, i < j\}$ is a $n_{edges} \times T$ matrix where $n_{edges} = 153$ is the number of ROI pairs and $T = 8$ is the number of phases. The SVD is performed over the non-centred matrix in order to obtain singular vectors in the general correlation matrix and not limit their directions in the normalized space of the average effect of morphine 70mg/kg.

As expected, the first non-centered moment (first mode) shows the same structure as the baseline matrix (Extended Data Fig. 8a). The second mode shows features of the morphine fingerprint (hippocampal shift from cortical to subcortical connectivity, high bilateral cortical correlation). As we performed a non-centered SVD, this mode can be used to compare similarity of any correlation matrix with either the baseline or the morphine fingerprint. Therefore, we define the FC index as the correlation between any correlation matrix and the second non-centered moment of the morphine 70mg/kg SVD. This FC index was normalized with 0% defined as its average at baseline, and 100% as the maximum cohort average correlation (morphine 70mg/kg, r = 0.61). Importantly, the FC index displays time- and dose -dependence (Extended Data Fig. 8b) similar to the morphine fingerprint shown on Fig. 2.

**Impact of animal mobility on FC and CBV**

As reported previously[5], in BALB/cJ mice opioid injection did not lead to augmented locomotor activity (Fig. 5a). However, morphine-induced transient CBV elevation and increase in the FC index are still present, albeit displaying distinct temporal dynamics and amplitude than in C57BL/6J mice (Fig. 5).

**Cross-correlation analysis**

To study the latency structure of the opioid fingerprint, the temporal cross-correlation between voxel and seed time series is computed, as shown on Fig. 3c of the main manuscript. We illustrate here the method of our lag analysis (Extended Data Fig. 3) and show Supplementary Videos showing how stationary and propagatory temporal cross-correlation patterns are massively and significantly reshaped by morphine injection.

**MOP in vivo phosphorylation assay**

HA-MOP knock-in mice[6] were either treated with agonists or received no treatment. Mice were anesthetized with isoflurane, killed by cervical dislocation, and brains were quickly dissected, excluding the cerebellum. Brain samples were immediately frozen in liquid nitrogen. Brains were transferred to ice-cold detergent buffer (50 mM Tris-HCl, pH 7.4, 150 mM NaCl, 5 mM EDTA, 1% Nonidet P-40, 0.5% sodium deoxycholate, 0.1% sodium dodecyl sulfate (SDS), containing protease and phosphatase inhibitors), homogenized, and centrifuged at 14,000 × g for 30 min at 4°C. The supernatant was then precipitated with Pierce™ Anti-HA Magnetic Beads (Thermo Fisher Scientific, Germany) for 60 min at 4°C. Afterwards the receptor-beads-conjugates were separated from the supernatant using a special magnetic device (DynaMag™-2, life technologies) and washed 3 times. Proteins were eluted from the beads with SDS-sample buffer for 25 min at 43°C and then resolved on 8% SDS-polyacrylamide gels. After electroblotting, membranes were incubated with the rabbit polyclonal phosphosite-specific MOP antibodies against pT370-MOP (7TM0319B), pS375-MOP (7TM0319C), pT376-MOP (7TM0319D), and pT379-MOP (7TM0319E) (7TM Antibodies GmbH, Jena, Germany)[6], followed by detection using a chemiluminescence detection system. Blots were subsequently stripped and incubated again with the phosphorylation-independent antibodies rabbit monoclonal anti-HA antibody (Cell Signaling, Frankfurt, Germany) and/or anti-MOP antibody {UMB-3} (Epitomics, Burlingame, CA) to confirm equal loading of the gels[6]. Films exposed in the linear range were then densitized using ImageJ 1.37v.

**Hot-plate test**

Opiate effects on paw withdrawal latencies were assessed as the time to response (licking or flicking fore or hind paw(s)) after placement on a hot-plate maintained at 56°C (Ugo Basile SRL, IT)[7]. To avoid tissue damage, we used a 30 s cut-off. The hot-plate test was carried out 15 min (fentanyl) or 30 min (morphine, methadone and buprenorphine) after drug administration and expressed as percent maximum possible effect (% MPE), calculated as follows: 100 × [(drug response latency − basal response latency)/(30 s − basal response latency)]. For time courses antinociception was determined at various time points after treatment. All calculations were performed using GraphPad Prism 6 software (GraphPad Software, Inc., San Diego, CA).

Acute paradigms for hot-plate test

For acute analgesia testing in the hot plate test, dose-response curves were generated after repeated subcutaneous administration of cumulative doses of fentanyl, morphine, methadone or buprenorphine. Mice were injected at 15 min intervals with 0.02, 0.03, 0.05, 0.1 and 0.1 mg/kg fentanyl to yield final cumulative doses of 0.02, 0.5, 0.1, 0.2 and 0.3 mg/kg fentanyl, and latencies were measured 15 min after drug administration, immediately followed by additional drug except for the last dose. For morphine dose-responses, mice were injected at 30 min intervals with 3.75, 3.75, 15 and 30 mg/kg morphine resulting in final cumulative doses of 3.75, 7.5, 22.5 and 52.5 mg/kg morphine. Hot plate latencies were always measured 30 min after morphine administration, immediately followed by the next dose. For methadone dose-responses, mice were injected at 30 min intervals with 2.5, 2.5, 2.5, 2.5 and 5 mg/kg methadone resulting in final cumulative doses of 2.5, 5, 7.5, 10 and 15 mg/kg methadone. For buprenorphine dose-responses, mice were injected at 30 min intervals with 0.1, 0.2, 0.7, and 2 mg/kg buprenorphine resulting in final cumulative doses of 0.1, 0.3, 1 and 3 mg/kg buprenorphine. For morphine dose-responses for BALB/cJ mice, mice were injected at 30 min intervals with 10, 20 and 40 mg/kg morphine resulting in final cumulative doses of 10, 30, and 70 mg/kg morphine. All dose-response curves generated using repeated cumulative dosing regimen are shown in Extended Data Fig. 6. Cumulative doses-response curves on wildtype (JAX™ C57BL/6J) mice after morphine and fentanyl treatment were already previously published in Kliewer *et al.* (2019) (Extended Data Fig. 6b)[7].

Tolerance paradigms for hot-plate testTo induce opioid tolerance for morphine and fentanyl, mice were implanted one day later (after the acute analgesia testing) with Alzet osmotic minipumps containing the same drug that was used in the acute cumulative dosing. Osmotic minipumps delivered total daily doses of 2 mg/kg fentanyl or 17 mg/kg morphine at a rate of 0.5 µl/h. Thus, in the tolerance paradigm dose-response curves from acute analgesia testing are depicted as day -1 and served as reference for the development of tolerance. On day 7, mice were again treated using a repeated cumulative dosing regimen with fentanyl (0.5, 0.5, 1, 2 mg/kg for WT) or morphine (22.5, 37.5, 60 mg/kg for WT), and hot plate response latencies were assessed at the same time points as on day −1. Cumulative doses-response curves on wildtype (JAX™ C57BL/6J) mice after chronic treatment of morphine and fentanyl were already previously published in Kliewer *et al.* (2019) (Extended Data Fig. 6b)[7].

**Respiratory depression**

Respiratory rates were recorded with a nose-out plethysmography system (Hugo Sachs Elektronik–Harvard Apparatus GmbH, DE) and already previously published in Kliewer *et al.* (2019)[7]. Data were used for correlation of morphine-induced FC changes and behaviour (Fig. 6 and Extended Data Fig. 7).

**Direct quantification of FC edges dysconnectivity as a function of time and morphine doses**

Dose and time dependence of morphine fingerprint could be quantified at the edge level (connectivity between two ROIs) for critical parts of the connectome. We show in (Extended Data Fig. 11) a detail of (Fig. 2) for bilateral cortico-cortical, cortico-hippocampal, cortico-thalamic and hippocampo-thalamic connectivity measures under (a.) various doses of morphine at BP2 and SP5 (b.) for 70mg/kg across all 8 sub phases of the recording. Statistics are computed using Mann-Whitney U test with multiple comparison correction using Bonferroni criterion.

**Stability of FC index estimate**

As the FC index is based on uncentered SVD it is a model-free, data driven quantity. For control, we studied the FC index estimate dependence on the dataset used for PCA. (Extended Data Fig. 12)shows that with the exception of the lowest dose (morphine 10mg/kg) the dynamics of FC is unaffected. We can also observe that the FC index tends to overfit the datasets it is trained on. IN order to be consistent with the dose dependence we report, FC index based analyses use the estimate trained on the highest dose (70mg/kg) which best preserves this hierarchy.

**CBV does not correlate with FC index**

We argue from Figure 6 that CBV and mobility follow a fast transient latent process uncorrelated from another slower one underlying FC, analgesia and phosphorylation. This result is further detailed Extended Data Figure 7. We demonstrate here that the weak negative correlation (r=-0.5, p=0.0132 ) between the FC index and CBV observed in both figures does not contradict this interpretation. Conversely, this anti correlation is driven by a spurious effect related to our choice of doing the correlation analysis only with the post-injection fUS data. Indeed, for analgesia and phosphorylation extension to pre-injection can only be done virtually where the value is zero by construction. For the sake of the argumentation we included these values in Extended Data Figure 13. Inclusion of the baseline data points preserves the strong correlations between the FC index, analgesia and MOP activation, while the spurious correlation between FC index and CBV disappears. This mechanism can be explained by the fact that the seemingly slow decay of CBV or mobility following the injection is actually a peak of fast transient activity which precedes the slow increase of FC index or analgesia.

**Extended Data Table 1**

**Extended Data Table 1** The following table recapitulates the analysed data for fUS experiments.

| **Experiment/Treatment** | **Aim** | **Selection** | **Reason** |
|---|---|---|---|
| Saline reference | 50 | 47 | misregistration |
| Saline control | 10 | 10 | - |
| Morphine 10 mg/kg | 10 | 9 | misregistration |
| Morphine 20 mg/kg | 10 | 10 | - |
| Morphine 30 mg/kg | 10 | 8 | misregistration |
| Morphine 70 mg/kg | 10 | 10 | - |
| Naloxone 2 mg/kg and morphine 70mg/kg | 10 | 8 | aberrant baseline |
| Fentanyl 0. 1 mg/kg | 10 | 10 | - |
| Fentanyl 0.2 mg/kg | 10 | 9 | misregistration |
| Fentanyl 0.25 mg/kg | 10 | 10 | - |
| Fentanyl 0.3 mg/kg | 10 | 10 | - |
| Buprenorphine 3 mg/kg | 10 | 9 | misregistration |
| Methadone 10 mg/kg | 10 | 10 | - |
| Morphine chronic 70 mg/kg | 40 | 40 | - |
| Fentanyl chronic 0.3 mg/kg | 40 | 40 | - |
| Saline BALB/cJ | 12 | 12 | - |
| Morphine 70 mg/kg BALB/cJ | 12 | 12 | - |
| Saline MOP$^{-/-}$ | 11 | 10 | death |
| Morphine 70 mg/kg MOP$^{-/-}$ | 11 | 10 | death |

**Extended Data Figure 1**

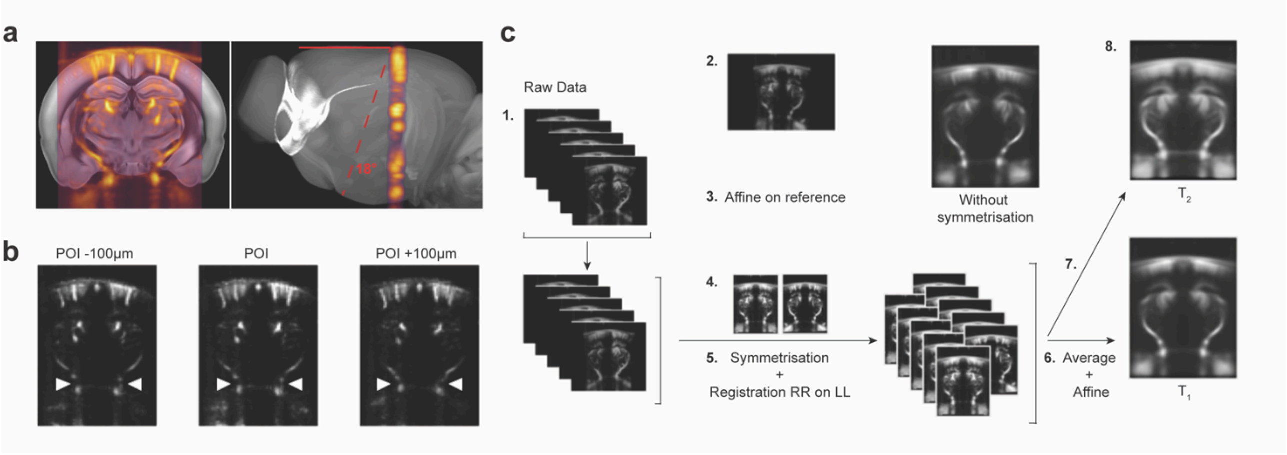


**Extended Data Figure 1 Probe positioning and slice registration. a,** Illustration of the imaged field of view in the implanted mouse (red line represents the coronal plane in the CCFv3). **b,** The plane of Interest (POI) can be identified with a 100 µm uncertainty by identifying the branching point between internal carotid and anterior choroidal artery. **c,** Template generation process as explained in the text (supplementary methods).

**Extended Data Figure 2**

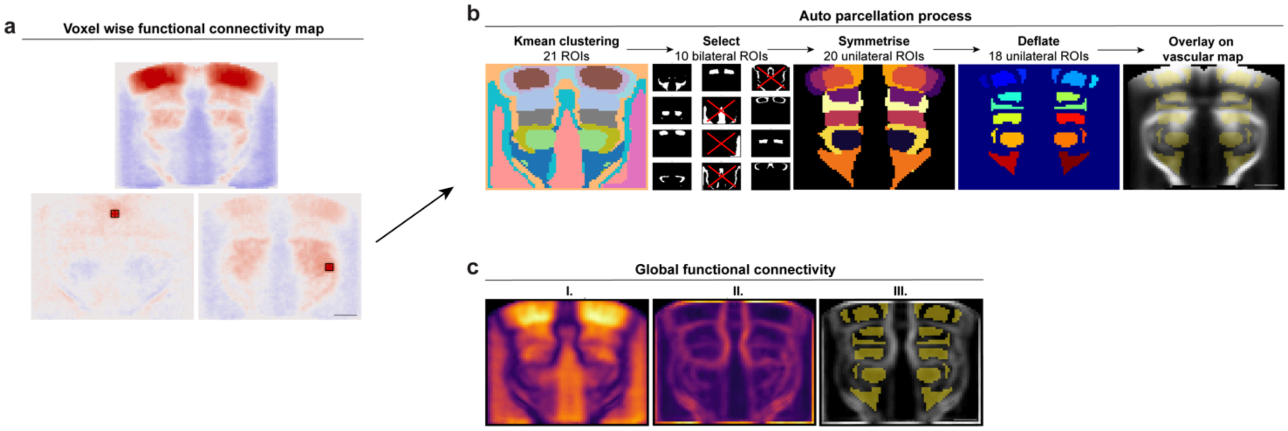


**Extended Data Figure 2 Automatic parcellation. a,** Two different example positions for a small 9 voxel seed (red square yield significantly different functional connectivity maps). **b,** From left to right: k-means clustering to identify voxel clusters displaying similar connectivity patterns; clusters adjacent to high-noise regions (mid-line and image boundaries) are discarded; remaining clusters are symmetrized to match the template symmetry; ROIs are deflated to mitigate misregistration issues. **c,** (I.) To validate the relevance of the automated parcellation, a global connectivity map is computed. (II.) The Laplacian of this map represents FC territory boundaries. (III.) ROIs are inside the boundary of this Laplacian map. Overlaying the parcellation with the Allen Brain Institute parcellation shows the anatomical relevance of this functional data driven approach.

**Extended Data Figure 3**

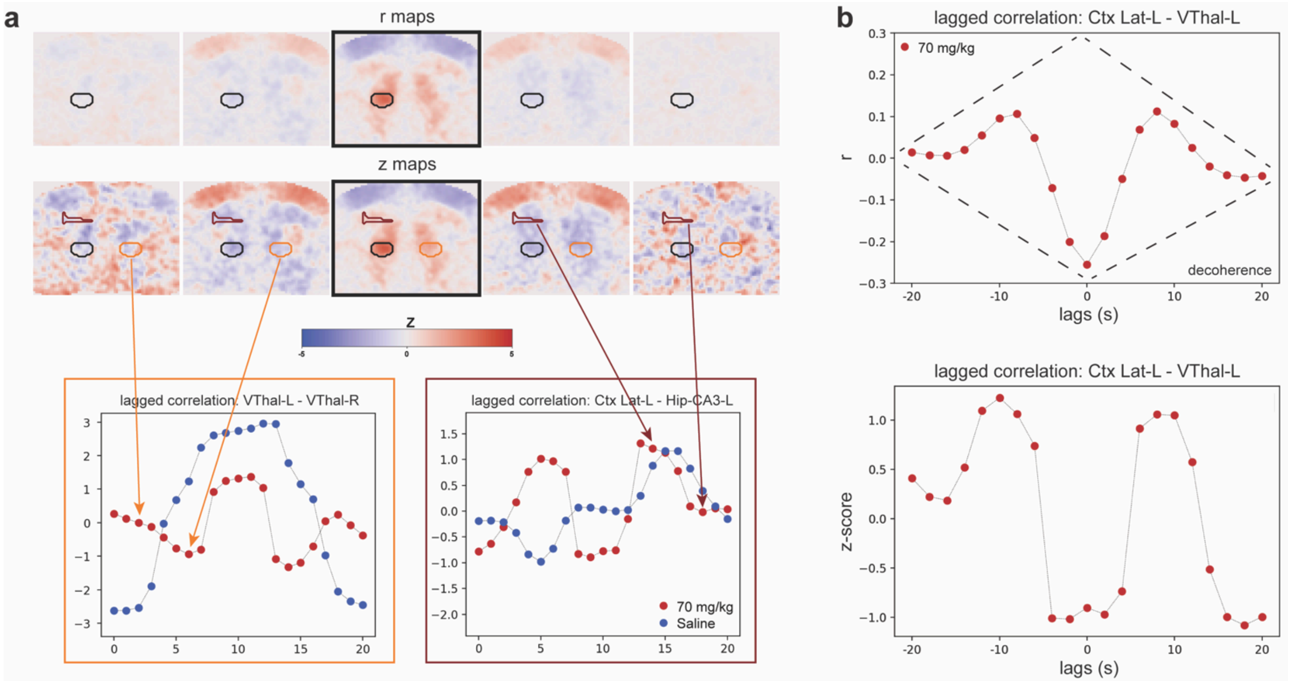


**Extended Data Figure 3 Example of lag analysis. a,** Illustration of lagged correlation time series generation, Top: cross-correlation r maps (blue-red [-5, 5]). Middle: each map is normalized to extract the z score, Bottom: the lagged correlation is computed as the average value of the lagged z scored SBM from a source seed (ventral thalamus in this case) to a target seed (orange: VThal, purple: Hip-CA3). The higher frequency (~20s period) oscillation induced by the morphine in the correlation pattern can be seen both for bilateral connectivity, but also inter regional one. **b,** Frame-wise transformation of r-scores to z-scores corrects for temporal decoherence.

**Extended Data Figure 4**

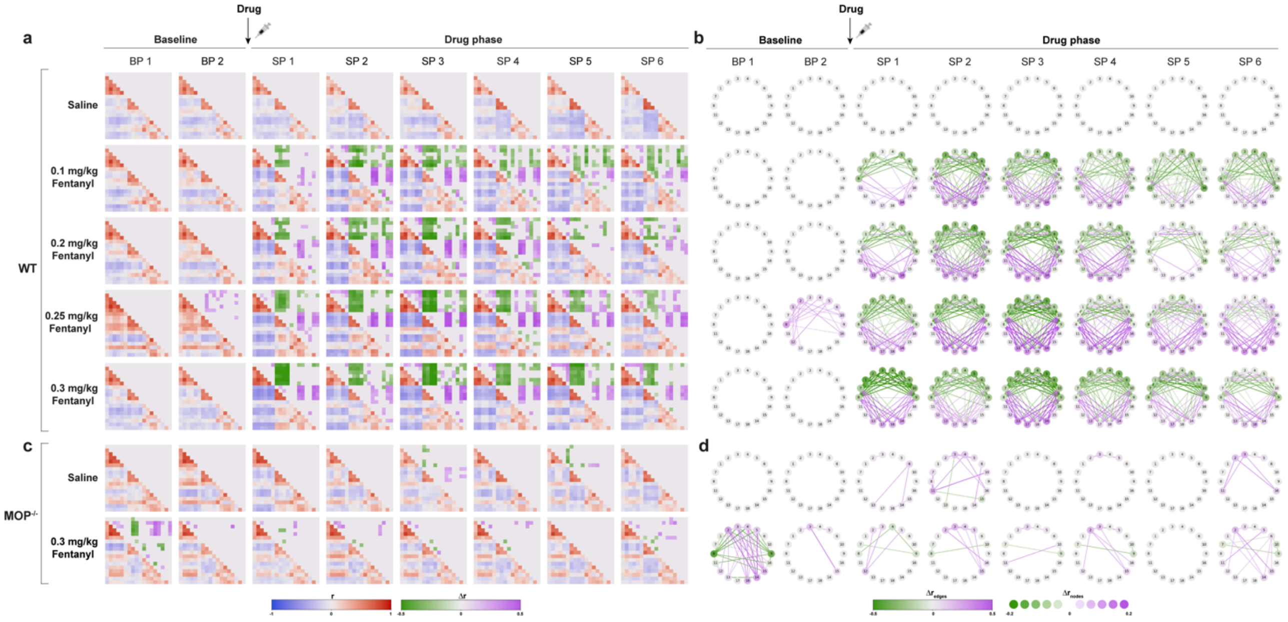


**Extended Data Figure 4 Fentanyl Dose response. a-c**, Pharmacodynamics and dose-dependence of the fentanyl FC pattern in **a, b,** wildtype (C57BL/6J) or **c, d,** MOP knockout mice (MOP$^{-/-}$) (n = 9 – 10). Statistical significance of the fentanyl FC pattern by comparing drug phase to saline injection. Correlation matrix in blue-red ([−1, 1]); significance matrix in green-violet ([−0.5, 0.5]); significance correlation graph in green-violet (edges [−0.5, 0.5]; nodes [−0.2, 0.2]). As compared to morphine (Fig. 2), the fentanyl FC fingerprint shows accelerated dynamics and effect size, matching the pharmacodynamics of fentanyl-induced analgesia (Suppl. Fig. 6).

**Extended Data Figure 5**

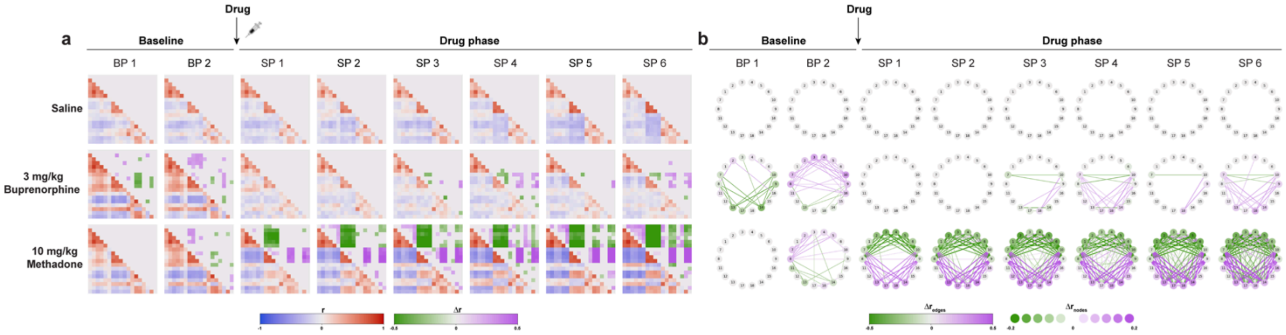


**Extended Data Figure 5 Pharmacokinetics of Buprenorphine and Methadone. a-b**, Pharmacodynamics and dose-dependence of the buprenorphine and methadone FC pattern in wildtype (C57BL/6J) (n = 9 – 10). Statistical significance of the drug FC pattern by comparing drug phase to saline injection. Correlation matrix in blue-red ([−1, 1]); significance matrix in green-violet ([−0.5, 0.5]); significance correlation graph in green-violet (edges [−0.5, 0.5]; nodes [−0.2, 0.2]). As compared to morphine (Fig. 2), these FC fingerprint shows altered dynamics and effect size, matching the pharmacodynamics of buprenorphine- and methadone-induced analgesia (Suppl. Fig. 6).

**Extended Data Figure 6**

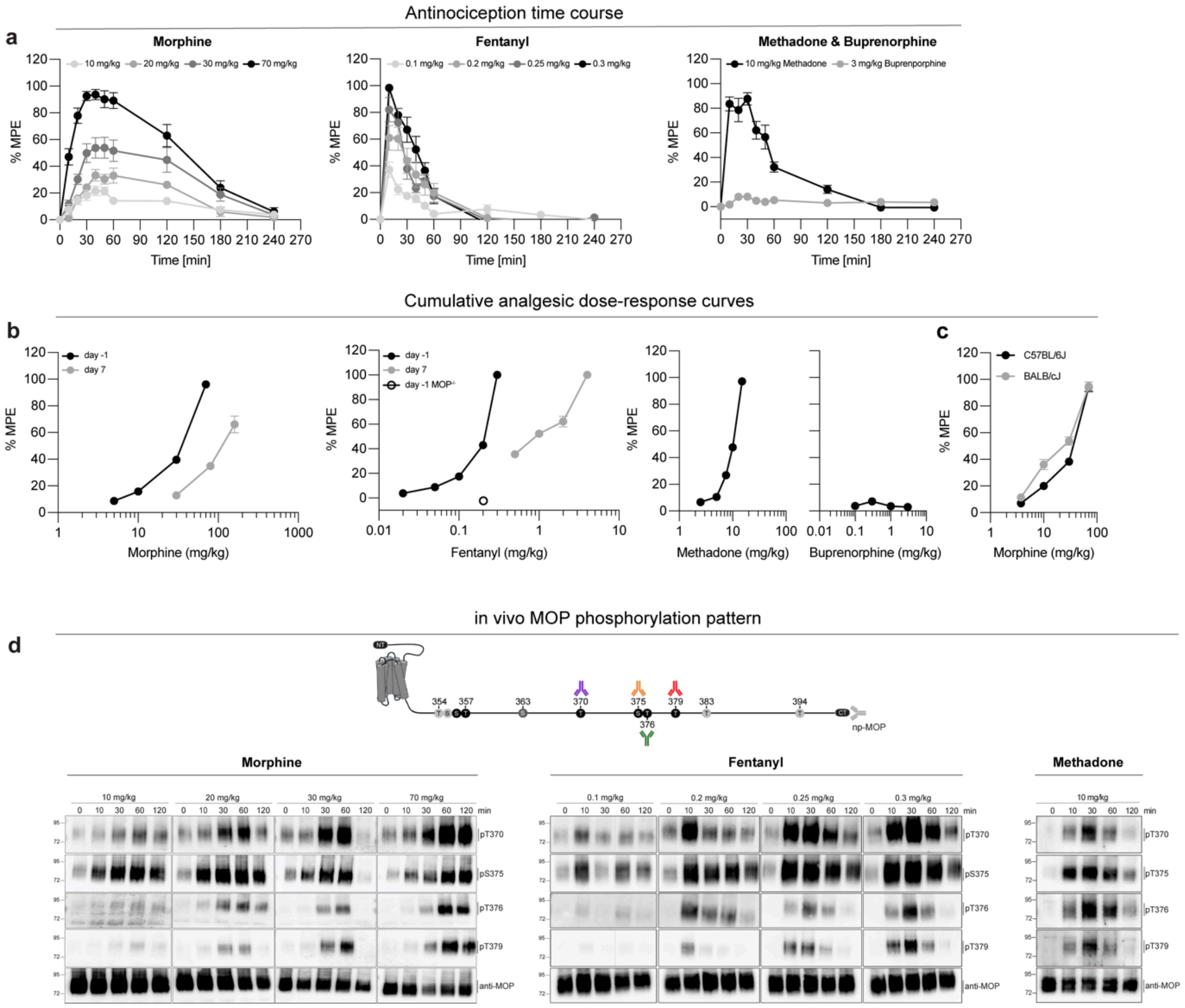


**Extended Data Figure 6 MOP effects on analgesia and phosphorylation. a,** Analgesic time course of acutely administered morphine, fentanyl, methadone or buprenorphine measured in the mouse hot-plate test was repeated after drug administration, respectively, for the indicated times (n = 6 – 17). **b-c,** Acute antinociceptive response measured in the mouse hot-plate test after fentanyl (15 min), morphine, methadone or buprenorphine (30 min) using a cumulative dosing regimen (n = 6 – 12). First two graphs: Cumulative analgesic dose-response curves before (day -1) and after (day 7) chronic treatment with osmotic pumps delivering morphine (17 mg/kg day$^{-1}$) or fentanyl (2 mg/kg day -1) (n = 9 – 12). **a-c,** Nociceptive latencies were defined by paw withdrawal and are reported as percent maximum possible effect (% MPE) with a 30-s cut-off. Data are the means ± s.e.m.. **d,** Schematic representation of the N-terminally tagged HA-MOP and the C-terminal tail with potential phosphorylation sites (black: antibody detectable phosphorylation sites; light grey: possible phosphorylation sites). HA-MOP knock-in mice were treated with different doses of morphine, fentanyl or methadone. Western blots of brain lysates from different treatment time points were immunoblotted with rabbit polyclonal phosphosite-specific MOP antibodies against pT370-MOP (7TM0319B), pS375-MOP (7TM0319C), pT376-MOP (7TM0319D), and pT379-MOP (7TM0319E). Blots were stripped and reprobed with rabbit monoclonal anti-HA antibody and/or anti-MOP antibody {UMB-3} to control for equal loading. Positions of molecular mass markers are indicated on the left (in kDa). Blots are representative of one to two independent experiments.

#### Extended Data Figure 7

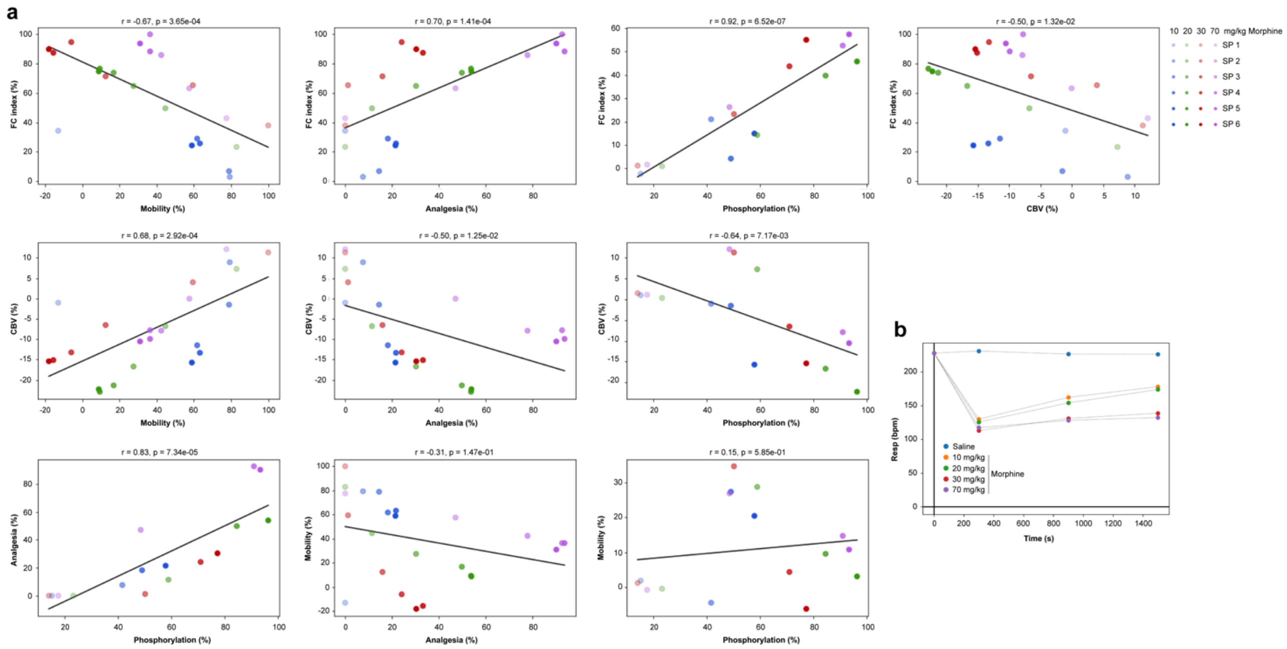

**Extended Data Figure 7 Detailed correlation patterns between the measured parameters. a,** Correlation between various parameters (FC, rCBV, mobility, analgesia, phosphorylation) following injection of morphine. each point represents a cohort average. Colors encode the injected dose (blue: 10 mg/kg, green: 20 mg/kg, red: 30 mg/kg and purple: 70 mg/kg) and time after injection (transparency increases with time). Pearson correlation coefficients and associated p-values. **b,** Average respiratory rate following injection of saline (blue) or increasing doses of morphine (orange: 10 mg/kg, green: 20 mg/kg, red: 30 mg/kg and purple: 70 mg/kg).

**Extended Data Figure 8**

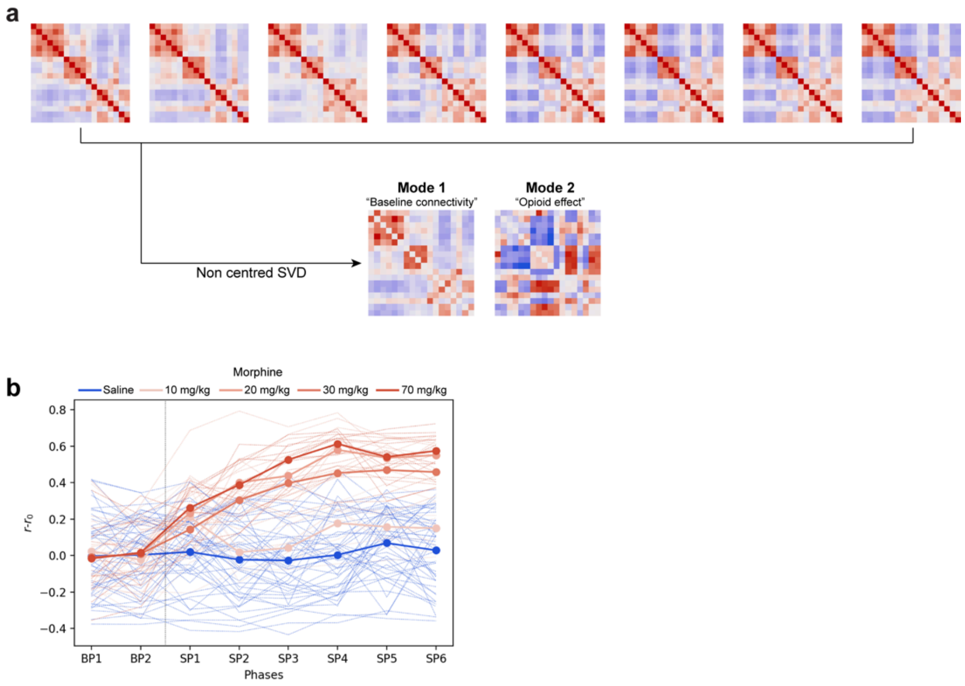


**Extended Data Figure 8 Illustration of the functional connectivity index. a,** The average FC matrices across all 8 phases of the morphine 70 mg/kg injection are computed. Using SVD on non-centered data, two modes are computed: the first mode represents the basal intrinsic connectome, the second mode shows morphine fingerprint specific features. **b,** When we compute the FC index after saline injection (blue) or different dose of morphine (red scale), the time and dose dependence of the fingerprint is well represented by the index.

**Extended Data Figure 9**

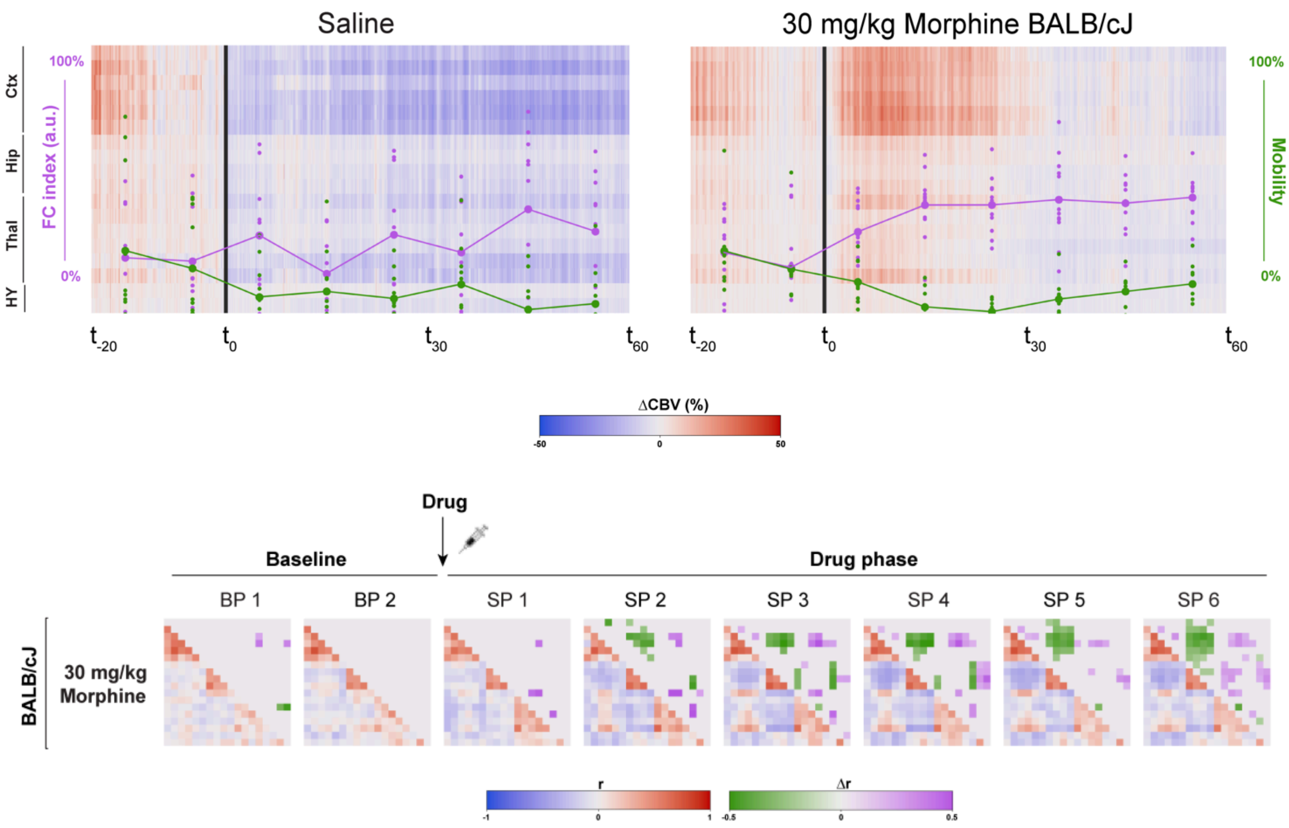


**Extended Data Figure 9 Impact of animal mobility on FC and CBV.** Top: Superimposition of functional connectivity changes (FC index in violet) and averaged mobility (green) on top of regional CBV change in blue-red ([−50, 50]) after injection of saline and morphine injection in wildtype (C57BL/6J) mice (n = 10) or BALB/cJ mice (n = 12), respectively. Bottom: Average and significance connectivity matrices following injection of morphine (30mg/kg) in BALBc mice showing the presence of the opioid fingerprint despite the absence of hyper locomotion.

**Extended Data Figure 10**

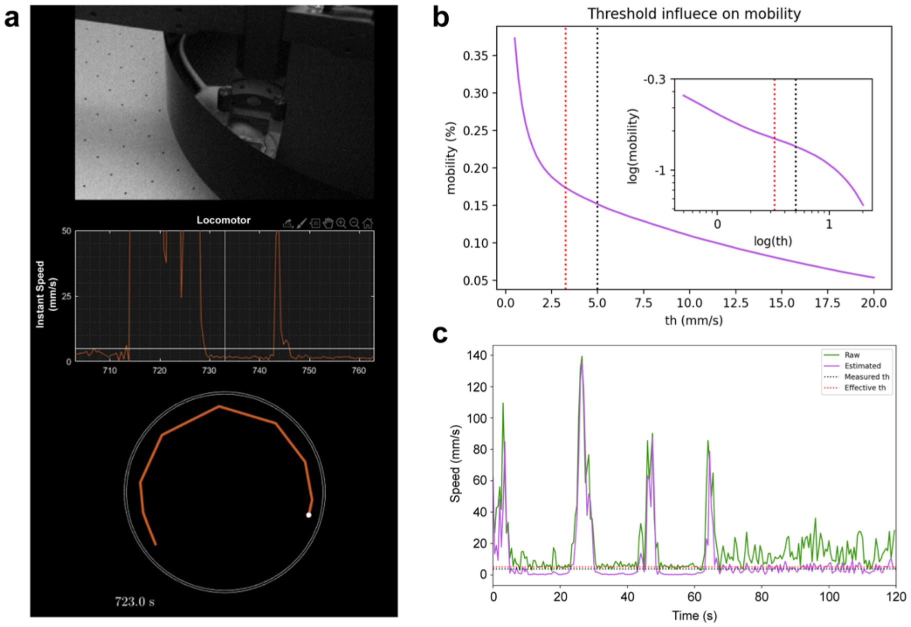


**Extended Data Figure 10 Mouse tracking during acquisitions. a,** The head-fixed animal (top) is tracked inside the Mobile HomeCage®, by directly measuring cage deplacement speed (middle) and translated cage position (bottom) with high precision. **b,** We identified two regimes (separated by the red dotted line) of movement depending on the estimated instant speed of the animal, corresponding to resting and movement periods. For the selection of genuine resting periods, we used a conservative criterion (black dotted line) in the linear domain to ensure robustness. **c,** We added supplementary robustness to the motion detection by re-estimating the instant speed on resampled position (purple) instead of directly using the resampled instant speed (green).

**Extended Data Figure 11**

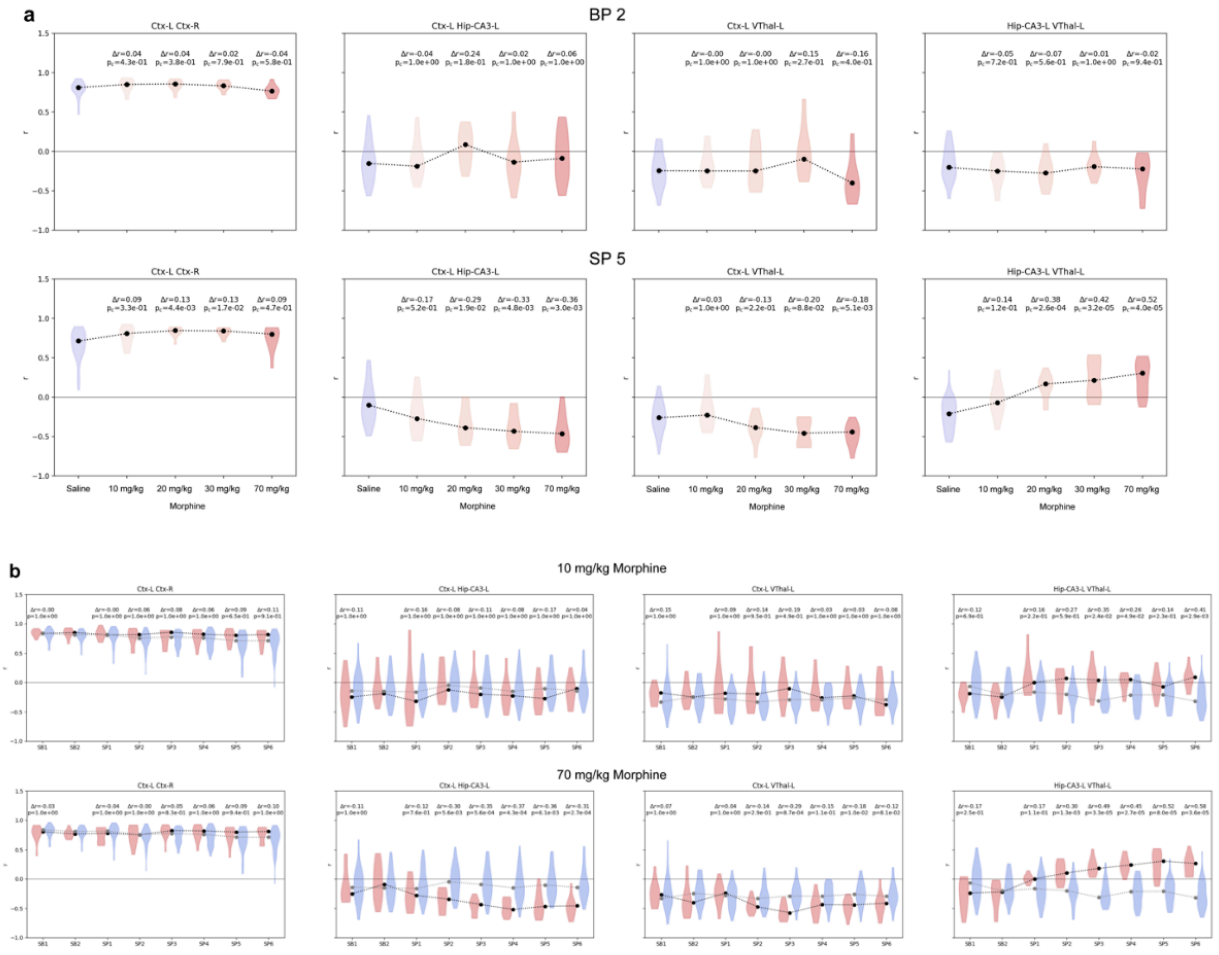


**Extended Data Figure 11 Edge level quantitative measure of morphine induced dysconnectivity.** Dose- and time- dependent FC changes following morphine treatment. a. : distribution and median of the Pearson's correlation coefficients between key ROI pairs of our network before (BP2: -10-0 mins) and after (SP5: +40-50mins) injection of saline or different doses (10, 20, 30, 70mg/kg) of morphine. b.: Temporal dynamics of FC in key ROI pairs of our network before and after injections of respectively 10mg/kg and 70mg/kg of morphine. Statistics are computed after Fisher transform (Mann-Whitney U test, corrected for multiple comparison with Bonferroni) and display is transformed back to r space for better readability.

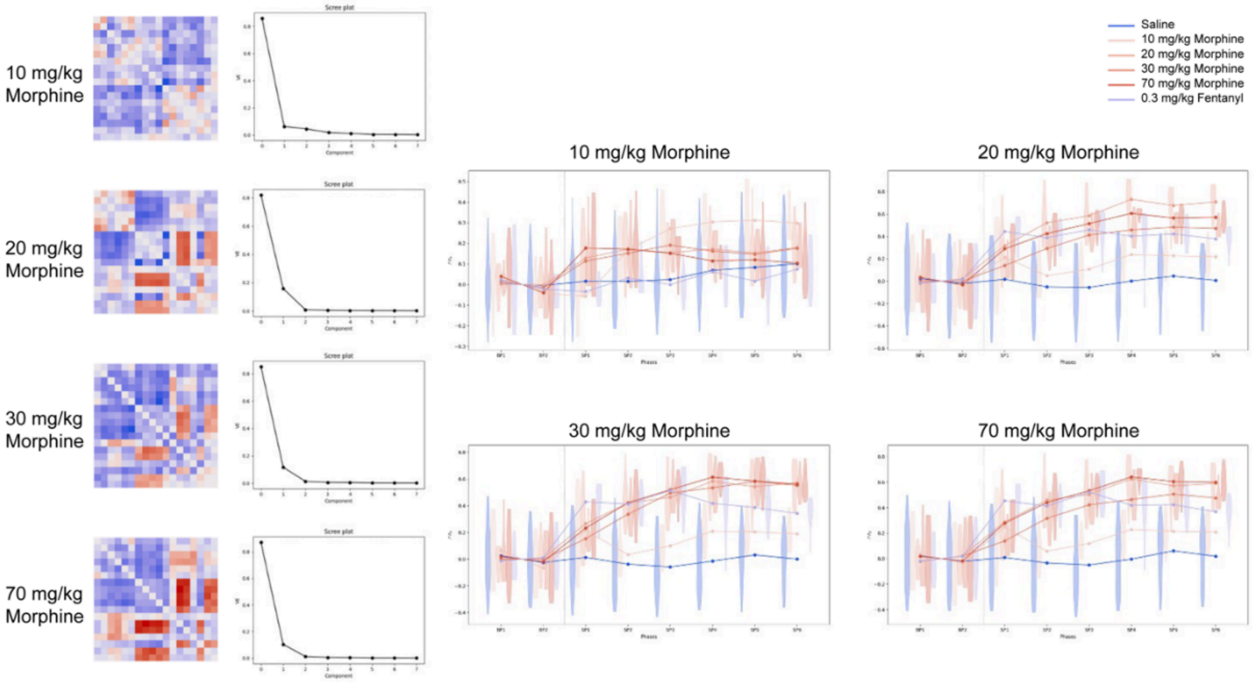


**Extended Data Figure 12 FC index stability.**

FC index trained for different doses of morphine. Left: spatial vector of the second component of the uncentred SVD which defines the FC index. With the exception of the 10mg/kg dose, main features of the "canonical fingerprint" topography are preserved. Middle: Scree plot of variance explained (VE) by the different components. Right: Temporal dynamics of the FC index for the different morphine doses follows the expected time- and dose-dependence with a slight overfitting towards the dataset used for training.

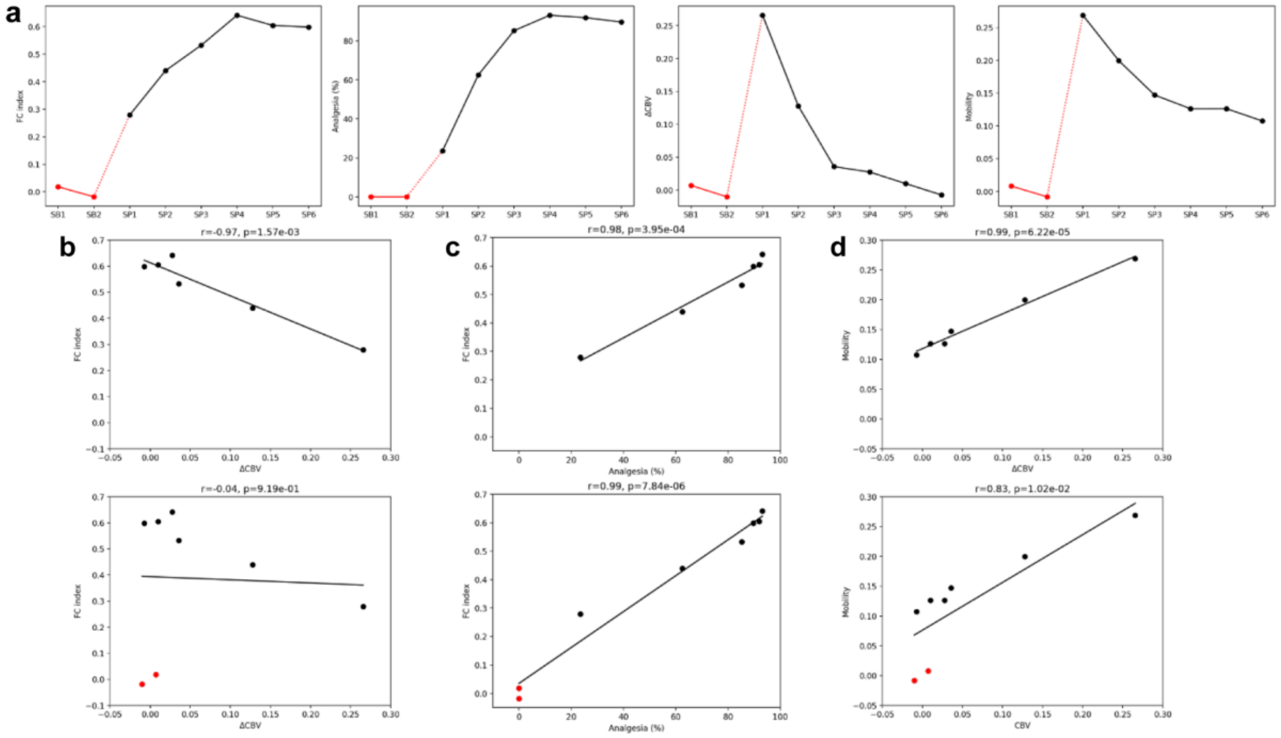


**Extended Data Figure 13 Spurious correlations in absence of baseline.**
Selecting data points only after injection can result in spurious positive correlation. a) timeseries from left to right: FC, analgesia, CBV and mobility show different influence of baseline (red dots) on their profiles. While considering only post injection (black) a monotonous increase (FC, analgesia) and decrease during post injection phase can induce a spurious correlation (b top graph). But, when baseline points are added, the correlation is lost (b bottom graph). Conversely, for both FC and analgesia (c) as well as CBV and mobility the correlation is preserved due to true linear dependence.

**Extended Data Video**

Each video represents one of the panels on Fig 3b-c with a lag of 0.5s between frames, with seeds in the lateral cortex (ROI 1), hippocampal CA3 (ROI 8), dorsal thalamus (ROI 11) and ventral thalamus (ROI 13). Right and middle panels show z-score maps (saturated at 5 standard deviations) at SP 5 after saline or 70 mg/kg morphine injection, respectively. The right panel show the voxels which present significant differences between the morphine and saline conditions in green-purple ($\Delta$r [-0.5, 0.5]).
